# A Bayesian Nonparametric Model for Inferring Subclonal Populations from Structured DNA Sequencing Data

**DOI:** 10.1101/2020.11.10.330183

**Authors:** Shai He, Aaron Schein, Vishal Sarsani, Patrick Flaherty

## Abstract

There are distinguishing features or “hallmarks” of cancer that are found across tumors, individuals, and types of cancer, and these hallmarks can be driven by specific genetic mutations. Yet, within a single tumor there is often extensive genetic heterogeneity as evidenced by single-cell and bulk DNA sequencing data. The goal of this work is to jointly infer the underlying genotypes of tumor subpopulations and the distribution of those subpopulations in individual tumors by integrating single-cell and bulk sequencing data. Understanding the genetic composition of the tumor at the time of treatment is important in the personalized design of targeted therapeutic combinations and monitoring for possible recurrence after treatment.

We propose a hierarchical Dirichlet process mixture model that incorporates the correlation structure induced by a structured sampling arrangement and we show that this model improves the quality of inference. We develop a representation of the hierarchical Dirichlet process prior as a Gamma-Poisson hierarchy and we use this representation to derive a fast Gibbs sampling inference algorithm using the augment-and-marginalize method. Experiments with simulation data show that our model outperforms standard numerical and statistical methods for decomposing admixed count data. Analyses of real acute lymphoblastic leukemia cancer sequencing dataset shows that our model improves upon state-of-the-art bioinformatic methods. An interpretation of the results of our model on this real dataset reveals co-mutated loci across samples.

## 1. Introduction

Intratumor heterogeneity is a major obstacle for the diagnosis and treatment of cancer. Genetic mutations that arise as the tumor grows produce clonal subpopulations (Vogelstein and Kinzler, 2004; Martincorena and Campbell, 2015), and resection of a fraction, but not all, of the tumor can alter the tumor environment in ways that provide a selective advantage to a remaining tumor clonal subpopulation leading to recurrence (Predina et al., 2013). Genomic instability in tumor cells results in a tumor where no single clonal population dominates the population (Hanahan and Weinberg, 2011). As a result, at a given point in time in the tumor development process, the population of tumor cells is a mixture of multiple genetic subpopulations (Lee et al., 2015; Russnes et al., 2011). The genetic composition of the tumor at the time of treatment is a critical factor in the design of targeted therapeutic combinations (Kyrochristos et al., 2019).

Tumor clonal subpopulations are genetic subpopulations whose constituent cells have acquired selected clonal driver mutations as well as unselected passenger mutations (Stratton, Campbell and Futreal, 2009). Such subpopulations are not necessarily completely genetically homogeneous; rather, they have greater similarity to each other compared to tumor cells that are not in the subpopulation (Chowell et al., 2018). Subclonal populations are subpopulations that represent less than 10% of the total tumor (Loeb et al., 2019). Additionally, a given patient sample can contain both tumor cells and normal cells; the purity of the sample is the ratio of cancer cells to total cells in the sample (Aran, Sirota and Butte, 2015).

The existence of clonal subpopulations has been known for many years (Nowell, 1976). Several reviews have covered the maintenance of heterogeneity in cancer samples (Bonavia et al., 2011; Marusyk, Almendro and Polyak, 2012). Gerlinger et al. (2012) showed that biopsies from regionally distinct locations in a solid tumor have different genetic mutations. Alizadeh et al. (2015) reviewed efforts to build consensus on definitions around tumor heterogeneity and highlights how understanding heterogeneity can inform therapeutic options. While the significance of tumor heterogeneity in treatment efficacy has been established, rigorous statistical modeling of tumor heterogeneity presents many challenges (Andor et al., 2016; Beerenwinkel et al., 2015).

Next-generation sequencing (NGS) has enabled the potential for the identification of subclonal populations in heterogeneous tumors with targeted sequencing of bulk samples (Campbell et al., 2008). Experiments that use material from millions of cells (bulk samples) can capture broad changes, but risk providing an average measurement that is not representative of the genetic state of any individual cell (Navin, 2015; Kalisky and Quake, 2011; Gawad, Koh and Quake, 2016). Recent advances in single-cell DNA sequencing have enabled researchers to collect sequence data using material from only a single cell (Treutlein et al., 2014). While single-cell experiments can capture the genetic state of the individual cell, sampling enough cells to gain a representative sample of population is expensive. Therefore, there is a need to integrate information from both bulk and single-cell data to obtain a comprehensive understanding of subclonal populations in an individual tumor as well as across individuals.

To summarize, Figure 1 shows a prototypical example of a heterogeneous tumor. A solid tumor is composed of three clonal subpopulations shown as divisions and numbered 1, 2, and 3. Three samples are obtained from the solid tumor; two are bulk, regional samples (samples 1 and 3) and one is a single cell sample (sample 2). DNA sequencing data from these samples is represented by bars above each genomic locus where the height of the bar is proportional to the number of observations of the nucleobase at the locus and the color represents the proportion of the observations with a mutated base. Subpopulation 1 is characterized by mutations at genomic locations 3(A) and 6(C) and subpopulation 2 is characterized by mutations at genomic locations 5(G) and 6(T). These true subpopulation genotypes are unknown and are inferred through the sequencing data from the biological samples. Sample 1, a bulk sample, is collected in a way such that a fraction of the cells are from subpopulation 1 and a fraction are from subpopulation 2 resulting in a mixture of observations from both subpopulations. Additionally, sequencing errors or passenger mutations may introduce observations of nucleobases that are not part of any true subpopulation — for example, at genomic locus 1. Bulk samples typically have good coverage and depth across genomic locations as shown by the presence of observations (bars) at each genomic locus and the relatively high height of the bars. For sample 2, the single-cell sample from subpopulation 2, the coverage is sparse and the depth is low, but the sample has does contain data that is relevant to inferring the single underlying subpopulation genotype. Sample 3 is a bulk sample from subpopulation 2 with no heterogeneity. In this sample a passenger mutation at genomic location 2 obscured the true subpopulation (2) genotype. This work aims to resolve the subpopulation genotypes and the distribution of the subpopulations for each tumor using multiple tumors samples from multiple individuals.

**Fig 1:**
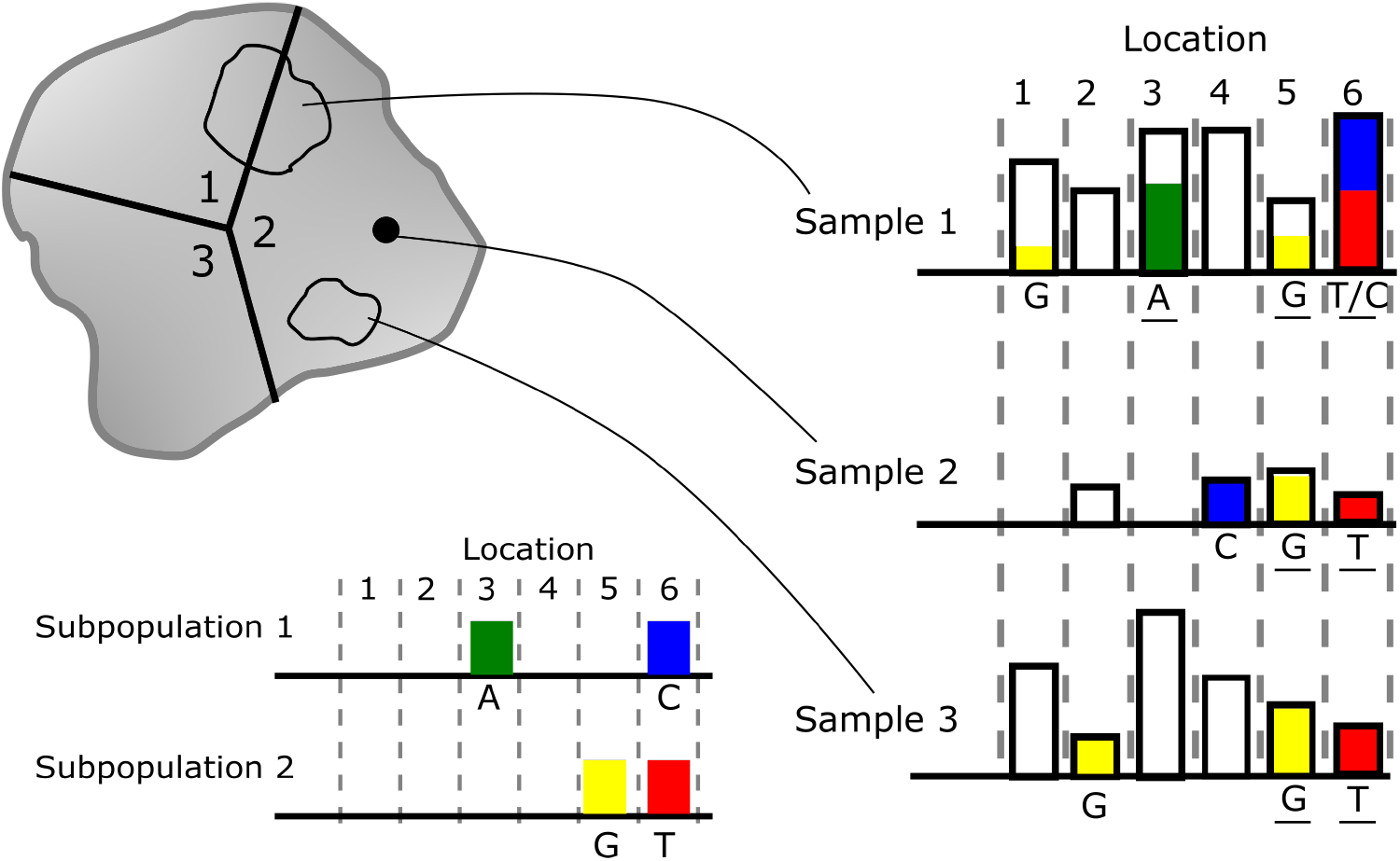
Prototypical example of tumor heterogeneity and clonal subpopulations.

### 1.1. Problem Setup

The fundamental unit of sampling in NGS data is the *read*. A read is a short DNA sequence of 100–400 nucleobases (bases) that maps to a specific location in a reference genome; a typical DNA sequencing run produces millions of such reads. We denote the observed DNA base in read *r* ∈ {1,…,*R_s_*} that maps to genomic location *l* ∈ {1,…,*L*} in sample *s* ∈ {1,…, *S*} as 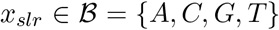. Since there are only four DNA bases, we have a 1-1 mapping from 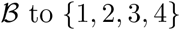 to {1,2,3,4}. When a genomic location has only two bases that are observed in a population the location is called *biallelic* and the sample space can be reduced to *x_slr_* ∈ {*A, a*}, where *A* is the major (most common) base and *a* is the minor (second most common) base. A read-count vector 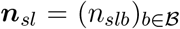 can be constructed by summing over the reads and conditioning on a genomic location, 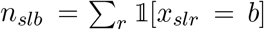. The *coverage* at a given location is 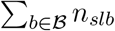 Goodwin, McPherson and McCombie (2016) present a comprehensive review of NGS, associated technologies, and summary statistics.

Single-cell sequencing data and bulk sequencing data differ in certain read-level statistics, but the fundamental observational unit for both is the read. DNA from single cells must be amplified by targeted amplification if a restricted region is of interest or whole genome amplification if the whole genome is of interest. The whole genome amplification process introduces false positives—apparent mutations that are not present in the original biological material and allelic dropout—heterogeneous alleles that appear homogeneous due to incomplete amplification of both alleles (Zafar et al., 2016). Both errors can be caused by founder effects due to early stage errors in polymerase chain reaction amplification. Additionally, singlecell sequencing data suffers from incomplete coverage of all loci and low sequencing depth (Zhang et al., 2019). In our problem setup each single-cell sample is treated as a single sample from the tumor.

Multiple NGS sequencing runs from an experiment are collected into a dataset, but the runs that comprise the dataset are rarely independent or identically distributed. In cancer sequencing datasets, there may be multiple individuals, each individual may have multiple solid tumors, and each solid tumor may have multiple biopsies. Data from model system experiments may have multiple genetic backgrounds and multiple environmental conditions. There may be multiple biological replicates and within each biological replicate there may be multiple technical replicates. A nested sampling structure produces samples that are correlated, and it is important to account for that correlation structure in the analysis of the data. In an experimental study, Paisley (2020) showed that a hierarchical Dirichlet process model performed better than a flat model when the number of samples at the lowest level of the sampling hierarchy was small. This data scenario is exactly the one we have with many NGS datasets.

### 1.2. Our contributions

The goal of this work is to jointly infer the underlying genotypes of tumor subpopulations and the distribution of those subpopulations for each tumor sample by making use of both single-cell and bulk sequencing data from multiple tumor samples in multiple individuals. In Section 2 we propose a Bayesian nonparametric hierarchical Dirichlet process mixture model for combining information from bulk and single-cell nextgeneration DNA sequencing data from multiple samples and from multiple individuals. This hierarchical Dirichlet process mixture model has tunable hyperparameters that control the a priori concentration of the subpopulation distribution for each sample; this hyperparameter can be estimated in an empirical Bayes setting or set directly when the concentration is known— for example, when the sample is from a single-cell. The hierarchical structure models the nested sampling structure in real NGS datasets that arises from drawing multiple bulk and single-cell biopsies from multiple individuals. Inference with our model provides estimates of the subpopulation genotypes and the distribution over subpopulations in each sample. In Section 3 we represent the model as a Gamma-Poisson hierarchical model and in Section 4 we derive a fast Gibbs sampling algorithm based on this representation using the augment-and-marginalize method. This representation and inference algorithm are generalizable to other models that make use of a hierarchical Dirichlet process prior and can be employed to derive a fast Gibbs sampler with analytical sampling steps for other models.

This work aims to identify subpopulations that contain both somatic and germline mutations. In any tumor sample, some “normal” cells are likely to be present — these contain germline mutations, but not somatic mutations. Therefore, the normal subpopulation is a valid latent subpopulation in the context of our problem setup. Furthermore, germline mutations that are shared across multiple individuals in a study population will be evident in inferred latent subpopulation genotypes. We compare our model to related work on modeling heterogeneous NGS data using simulation experiments (Section 5) and we analyze real NGS data from a acute lymphoblastic leukemia (Section 6). Statistical inference provides estimates of the subpopulation genotypes and the distribution of subpopulations in individual samples with rigorous Bayesian uncertainty estimates. Since our inference algorithms produce samples from the full posterior distribution, our methods allow for rigorous quantification of the uncertainty in our estimates.

### 1.3. Related Work

We briefly review related work in the area of Bayesian nonparametric modeling using the Dirichlet process and in the area of bioinformatic analysis of genetically heterogeneous samples.

#### 1.3.1. Hierarchical Dirichlet Process Mixture Models

In real data sets there is often structural information that can increase the utility of the data towards an inferential task. One of the most common pieces of structural information is the a priori similarity among related samples. As an example, suppose that we have a set of news articles and we are interested in drawing inferences about the topics in the articles. A naïve model might assume that all articles are independent samples; a more sophisticated model would incorporate information about the authorship—articles by the same author are a priori likely to be more similar to each other than to articles by different authors. In the Bayesian formalism, the hierarchical Dirichlet process enables one to incorporate such structural information in the inference process in a rigorous model-based way.

##### Dirichlet Process

The Dirichlet process, *G* ~ DP(*α*_0_,*G*_0_), is formally a measure on measures where *α*_0_ > 0 is the scaling parameter and *G*_0_ is the base measure (Ferguson, 1973). A constructive definition is the stickbreaking representation (Sethuraman, 1994)

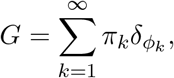

where

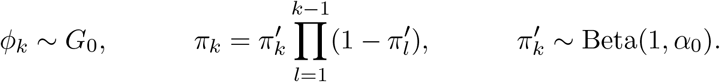

The sequence 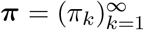 can be interpreted as a random probability measure on the positive integers and each integer is associated with a draw from the base measure. A second perspective of the Dirichlet process makes the clustering property more evident. The Chinese restaurant process (Aldous, 1985) describes a stochastic process where a draw *θ_i_* associates with a parameter *ϕ_k_* according to all of the previous pairs,

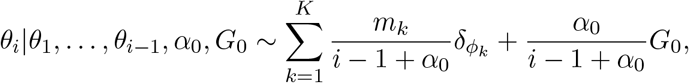

where *m_k_* is the count of *θ_i_*’s that are equal to *ϕ_k_*. The conditional distribution is a mixture distribution where the weights are determined by the previous draws and *α*_0_. If *α*_0_ is large, the mixture distribution is weighted towards new draws from the base measure, *G*_0_, and if *α*_0_ is small, it was weighted towards previously sampled values of *ϕ_k_*, where *ϕ*_1_,, *ϕ_K_* are the distinct values taken on by *θ*_1_,…,*θ*_*i*–1_. For this reason, *α*_0_ is called the *concentration* parameter of the Dirichlet process.

##### Dirichlet Process Mixture Model

The Dirichlet process is a natural nonparametric prior for models that need a probability measure. In particular, it is useful for mixture models because they employ a probability measure as a latent or unobserved variable. In parametric models, a natural prior for this latent variable is a Dirichlet distribution. By substituting a Dirichlet process, the number of mixture components scales with the size of the data set and in the asymptotic limit of the sample size, *n* → ∞, the number of components goes to infinity, *K* → ∞. The Dirichlet process mixture model can be written as

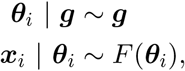

where *F*(***θ**_i_*) is the sampling distribution of the observed data, ***x**_i_*. The Dirichlet process mixture model can be construed as the infinite limit of a particular finite mixture model (Neal, 1992; Rasmussen, 2000; Green and Richardson, 2001; Ishwaran and Zarepour, 2002). The finite mixture model is

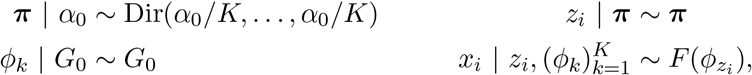

where *ϕ_k_* is the parameter for mixture component *k* drawn from prior distribution *G*_0_, and *z_i_* is an indicator of the mixture component. In the limit as *K* → ∞, this finite mixture model converges in distribution to the Dirichlet process mixture model (Ishwaran and Zarepour, 2002).

##### Hierarchical Dirichlet Process Mixture Model

The hierarchical Dirichlet process is a hierarchical extension of the Dirichlet process mixture model where the prior over the mixing Dirichlet process is itself drawn from a Dirichlet process. Given a base measure *H* and concentration parameter *γ*, the hierarchical Dirichlet process mixture model is

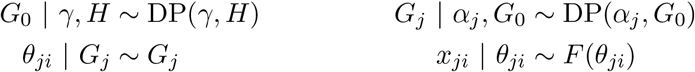

This hierarchical stacking can be extended in the direction of the prior.

#### 1.3.2. Bioinformatic models for clonal subpopulation inference

There are many methods for inferring the clonal genetic subpopulation structure from next-generation DNA sequencing data. A subset of these methods are based on a rigorous statistical model. We briefly review the most popular model-based bioinformatic methods for subpopulation structure inference. A more complete review of methods for subclonal inference is provided in Appendix A. PurityEst (Su et al., 2012) and PurBayes (Larson and Fridley, 2013) make use of paired tumor-normal samples. Roth et al. (2014) proposed a Dirichlet process mixture model for subpopulations called Pyclone. Phy-loWGS uses a Bayesian nonparametric model to reconstruct genotypes of the subpopulations from sequencing data (Deshwar et al., 2015). Bayclone uses an Indian buffet process prior over the genotypes for the subpopulations (Sengupta et al., 2015). Sciclone uses a hierarchical Bayesian mixture model to infer subclonal populations (Miller et al., 2014). CloneHD integrates information from copy number data, B-allele frequency, and somatic nucleotide variants to infer clonal subpopulations (Fischer et al., 2014). Cloe takes the innovative approach of incorporating a prior over phylogenetic trees (Marass et al., 2016). Treeclone is a nonparametric Bayesian model for reconstructing the clonal subpopulation phylogeny and inferring tumor heterogeneity (Zhou et al., 2019a).

##### Our work

Our work handles multiple subpopulation components in the tumor unlike paired tumor-normal methods — paired data is not required. Like Pyclone, we use a Dirichlet process prior over the samples. Our model uses a simpler prior over the subpopulation genotypes compared to Bayclone, and uses a hierarchical Dirichlet process prior over the samples instead of an Indian buffet process. This modeling choice enables us to focus on posterior inference for the subpopulation genotypes and the distribution over genotypes.It has been shown that while the posterior distribution of the Dirichlet process is consistent, inference on the number of components is not (Miller and Harrison, 2013, 2014). For this reason, we focus on the posterior distribution of subpopulation genotypes and the posterior distribution of subpopulations for each sample.

## 2. Hierarchical Dirichlet Process Mixture Probability Model

The full hierarchical Dirichlet process mixture model can be decomposed into the following components: the sampling model (Section 2.1), the hierarchical prior (Section 2.2), and the hyperparameters (Section 2.3). The full model and the complete posterior distribution is summarized in Section 2.4.

### 2.1 Sampling Model

The model presented here assumes biallelic variants with the major allele denoted by *A* and the minor allele denoted by *a*, but the model is easily adapted for a situation where *x_lsr_* records the observed DNA base {*A,C,G,T*}. The set of genotypes is denoted 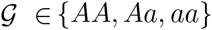 for a diploid genome and can equivalently be represented as 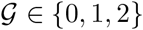. We assume a conditional categorical sampling model for *x_slr_*,

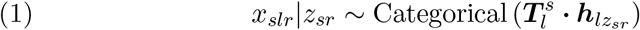

where

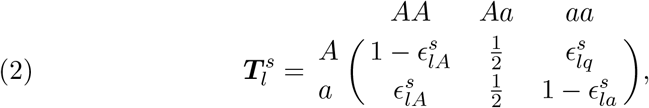

is the genotype-base transition matrix. This matrix is typically the product of a sequencing error model and can be specified for a particular location *l* and for a particular sample *s*. The hyperparameters 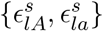 can be estimated from historical data on the sequencing error rate at location *l* and set distinctly for bulk sequencing samples or single-cell samples due to the dependency on the sample *s* = (*i,j*).

The genotype for subpopulation *k* at location *l* is 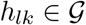. The categorical variable *h_lk_* can be equivalently represented by categorical indicator vector 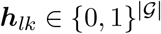. The conditional distribution of the genotype of subpopulation *k* at location *l* is

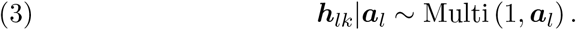

for all *l* = 1,…, *L* and *k* = 1,…, *K*. A simple independent prior for ***h**_lk_* using Hardy-Weinberg equilibrium can be used, 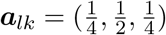, or the prior should be adjusted based on population frequency information.

The integer-valued variable *z_sr_* ∈ {1,2,…, *K*} indicates the genetic subpopulation that produced read *r* in sample *s*. It has a categorical conditional distribution

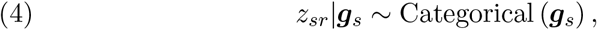

where ***g**_s_* is the distribution over subpopulations for sample *s*.

### 2.2. Hierarchical Prior

A key aspect of our model in the context of DNA sequencing datasets is a hierarchical Bayesian nonparametric prior on the distribution of subpopulations in sample *s*. Let sample *s* be generated by first drawing *individual i* = 1,…,*N* from a population and then drawing *biopsy j* = 1,…, *N_i_* within individual *i*. Therefore, the sample is *s* ∈ {(*i,j*) | *i* = 1,…, *N, j* = 1,…, *N*}. The following hierarchical Dirichlet process prior is used for modeling ***g**_s_* = ***g**_ij_*,

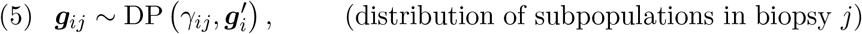

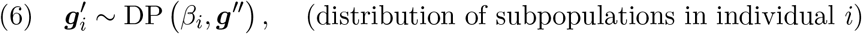

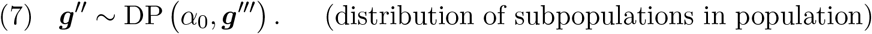

Here, ***g**_ij_* is the distribution over subpopulations in biopsy *j* from individual *i*, 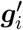 is the distribution over subpopulations in individual *i*, and ***g***″ is the distribution over subpopulations in the population from which the individuals are drawn. The top level prior measure ***g***′″ together with the concentration parameter *α*_0_ defines the prior over the population-level distribution of subpopulations. The products of inference in this model include the posterior distribution of these quantities.

### 2.3. Hyperparameters

The hyperparameters *α*_0_, *β_i_*, and *γ_ij_* are important for modeling single-cell and bulk sequencing experiments. If *γ_ij_* is set to a small value, then ***g***_*ij*_ is expected to be concentrated to one of the subpopulations. Therefore, if sample *s* = (*i, j*) is known to be from a single-cell, *γ_ij_* can be set to a small value to represent an expected concentration to a single subpopulation. If the sequenced sample is a bulk of cells or an entire solid tumor, *γ_ij_* can be set to a large value to represent an a-priori expectation of tumor heterogeneity. Hyperparameters *α*_0_ and *β_i_* represent prior information about subpopulation concentration at higher levels of the model. If *β_i_* is set to a small value, the distribution of subpopulation for individual i is concentrated on a small number of subpopulations. Since ***g***_*ij*_ is conditioned on 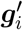, the individual concentration parameter, *β_i_*, influences the concentration of all of the biopsies within the individual. This hierarchical concentration in the model is congruent with the biological expectation that if a subpopulation is not present at the level of the individual, it would not emerge spontaneously in a sample from the individual. If *α*_0_ is set to a small value, the subpopulation distribution is, a-priori, concentrated at only a few subpopulations. The inclusion/exclusion criteria for the dataset can therefore influence the concentration of the entire population. If the dataset contains only a small subset of the entire population, for example a subset of triple-negative breast cancer patients in a clinical trial, it may be reasonable to set *α*_0_ to a small value. In the standard Bayesian paradigm, if the concentration parameter is not known a-priori, the associated parameters can be endowed with a Gamma distribution as in Escobar and West (1995).

### 2.4. Complete Hierarchical Dirichlet Process Model

By combining the sampling model and the hierarchical prior the complete hierarchical Dirichlet process model is

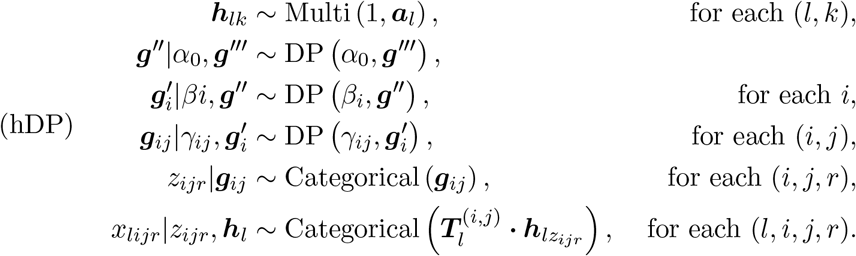

A graphical model representation of Model hDP is shown in Figure 2. Model hDP is conceptually compared with other common hierarchical models for factorizing count data in Appendix D.

**Fig 2:**
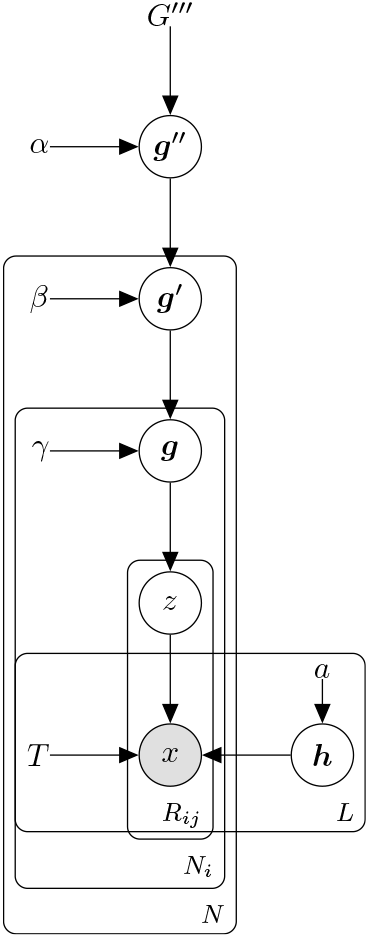
Graphical model representation of Model hDP.

The object of inference is the posterior distribution function for Model hDP:

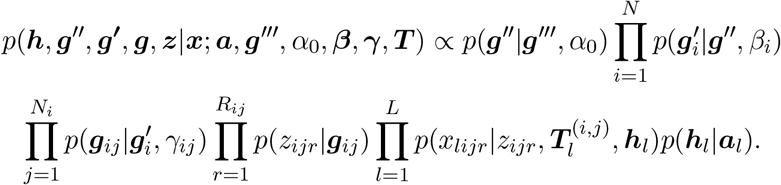

Next, we derive a Markov chain Monte Carlo (MCMC) inference algorithm to estimate this posterior distribution.

### 2.5. Inference Algorithm for Hierarchical Dirichlet Process Model

It is well-known that a Dirichlet distribution with parameter 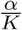 converges to a Dirichlet process as *K* → ∞ (Teh et al., 2006; Ishwaran and Zarepour, 2000). We employ this fact to derive an MCMC algorithm to draw samples from the posterior distribution. The details of the derivation of the truncated Dirichlet process inference algorithm are in Appendix B. The algorithm itself is shown in Algorithm 1.

## 3. Hierarchical Gamma-Poisson Probability Model

The inference algorithm for Model hDP employs Metropolis-Hastings steps that can be computationally expensive for large datasets. To address this issue, we reformulate the model as a hierarchical Gamma-Poisson model. This reformulation allows us to use the augment-and-marginalize method developed by Zhou et al. (2012) to derive a fast inference algorithm that uses only analytical sampling steps for updates.

### 3.1. Sampling Model

In the hierarchical Dirichlet process model, the observed data is the base for each read; in this Gamma-Poisson reformulation, the observed data is count of reads associated with each base. Let *Y_ijlb_* ∈ {0,1,2,…} be the read count of base 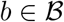 at location *l* ∈ {1,…, *L*} in biopsy *j* ∈ {1,…, *N_i_*} of individual *i* ∈ {1,…, *N*} (recall we have defined a sample as the pair *s* = (*i,j*)). We assume the read count has a conditional

**Algorithm 1:**
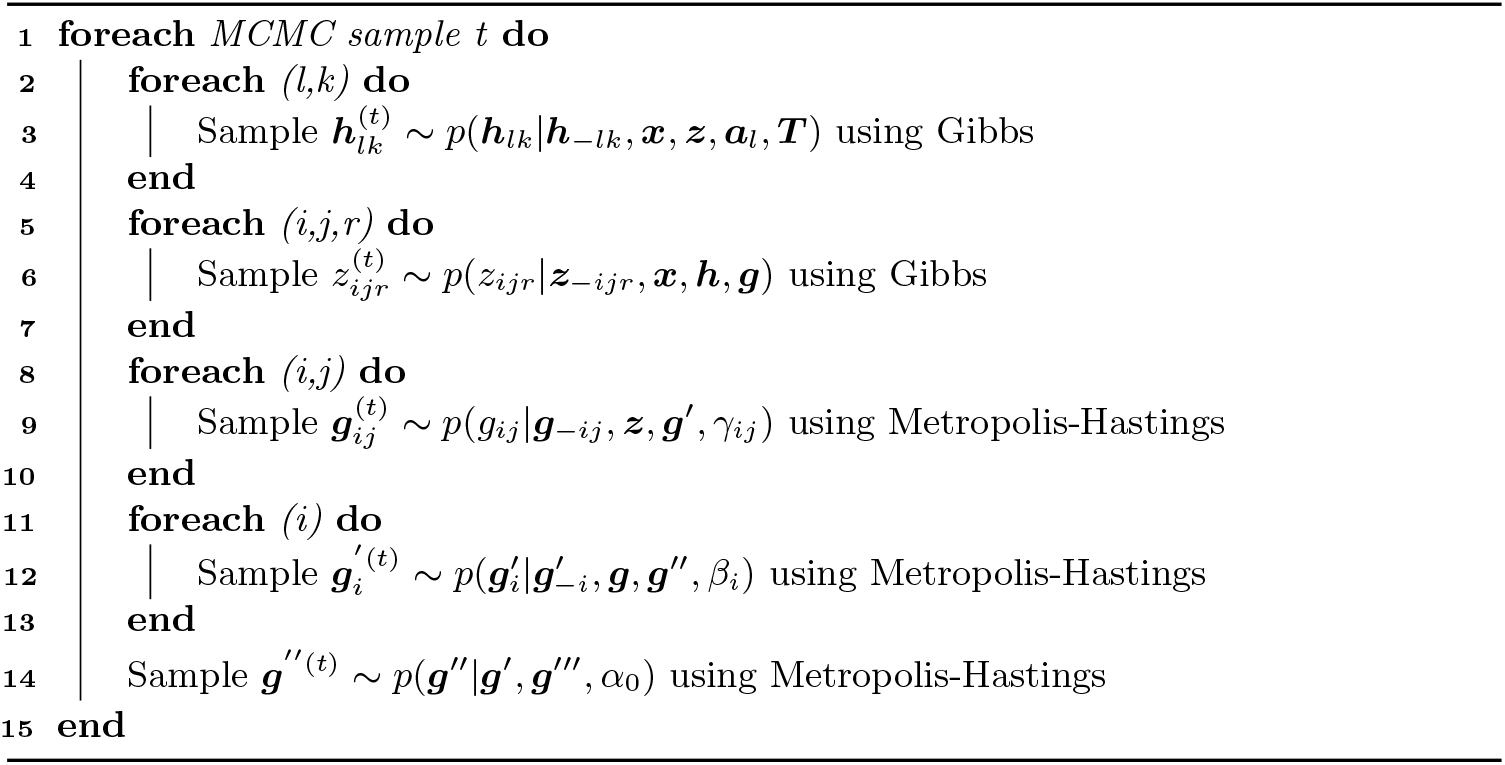
MCMC sampler for Model hDP

Poisson distribution,

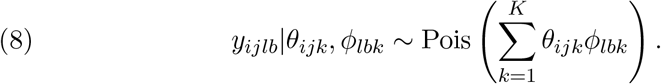

Note that while the conditional distribution is Poisson, the marginal distribution is negative binomial as shown in Equation (15). The rate parameter of the Poisson is a sum over *K* subpopulations where the summand is the product of two factors. The first factor *θ_ijk_* is the rate or propensity of subpopulation *k* in sample *s* = (*i,j*). The second factor is *ϕ_lbk_* = (***T***_*l*_ · ***h***_*lk*_)_*b*_ ∈ (0,1) and can be interpreted as the probability of base *b* in subpopulation *k* at location *l*. This representation requires the same genotype-nucleobase transition matrix across all samples.

### 3.2. Hierarchical Prior

We assume the following hierarchical gamma prior for propensity *θ_ijk_*:

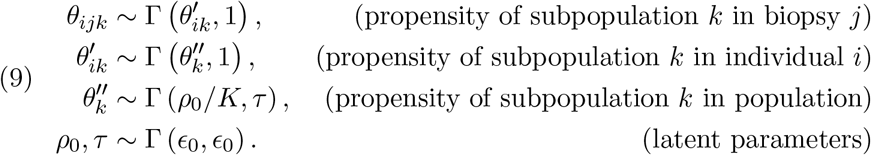

The form of the gamma distribution is Γ (*a, b*) where a is the shape parameter and b is the rate parameter. While the gamma distribution is the conjugateprior to its own rate parameter, this hierarchical prior is in a non-conjugate configurations since it chains through the shape parameter. Nevertheless, this construction yields closed-form complete conditional distributions via an auxiliary variable augment-and-marginalize update that is derived in Section 4.

### 3.3. Hyperparameters

The hyperparameter *ϵ*_0_ can be set to a small value for a diffuse prior over the parameters *ρ*_0_ and *τ*. If it is known that the entire data set is relatively concentrated on only one subpopulation *ϵ*_0_ can be set to a smaller value such as one. Or, if there is a priori information about the expected number of subpopulations, the shape and rate parameters can be adjusted accordingly. The distributions are not restricted to depend on a single hyperparameter.

### 3.4. Complete Gamma-Poisson Model

The complete Gamma-Poisson model is

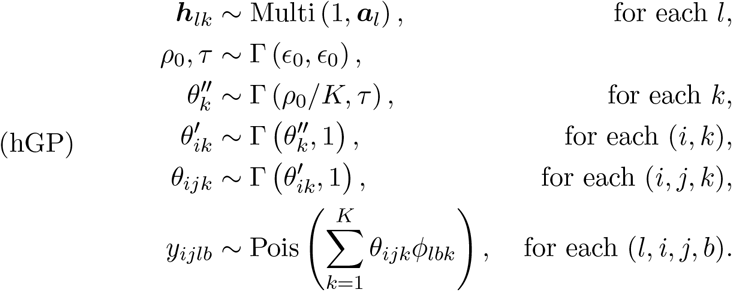

This model trades model flexibility for computational efficiency. The transition matrix ***T**_l_* is fixed for all samples and there is no tunable prior subpopulation concentration of each sample, but the computational efficiency of the resulting inference algorithm is significantly better than the hierarchical Dirichlet process mixture model and this model can be fit to much larger data sets. A complete graphical model representation of Model hGP is shown in Figure 3.

**Fig 3:**
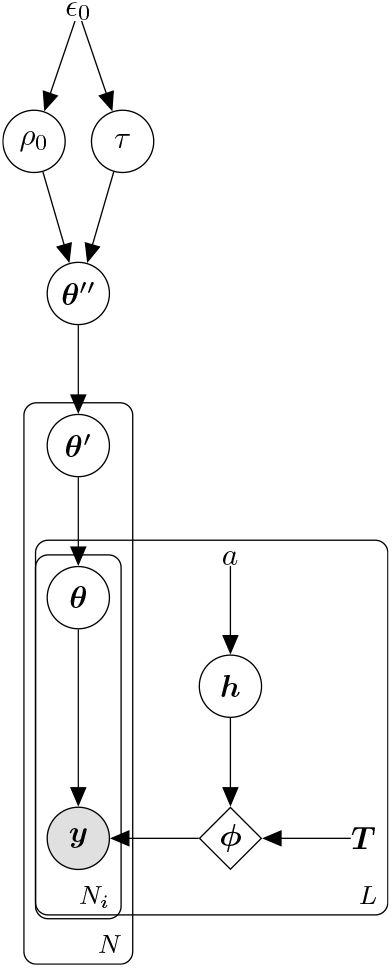
Graphical model representation of Model hGP.

The complete posterior distribution of the data under the Gamma-Poisson model is

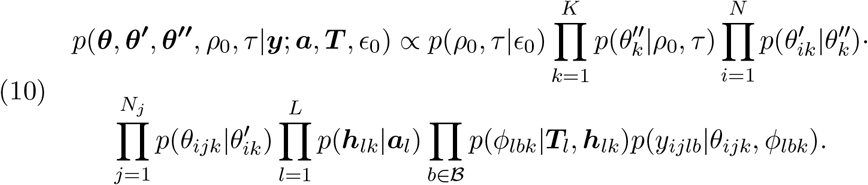

### 3.5. Interpretation as a Hierarchical Dirichlet Process Mixture Model

The hierarchical prior in Equation (9) can be interpreted in terms of a hierarchical Dirichlet process, similar to that given in Section 2.2. To see this, we appeal to (1) the relationship between the Gamma and Dirichlet distributions, and (2) the limiting form of the finite-dimensional Dirichlet distribution as the number of subpopulations goes to infinity.

A sample from a Dirichlet distribution can be obtained by normalizing a vector of independent gamma random variables with equal rate parameters but possibly different shape parameters. Suppose *θ_k_* ~ Γ(*a_k_*, 1) for *k* = 1,…,*K* are *K* independent Gamma random variables with shape parameters *a_k_*. We adopt dot (·) notation 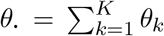 to denote sums. We denote proportion vector 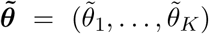 where 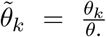. Lukacs (1955) showed that *θ*. and 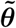 are then marginally (i.e., not conditional on *θ*_1_,…,*θ_K_*) independent; moreover, they are distributed as *θ*. ~ Γ(*a*., 1), where 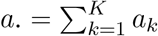, and 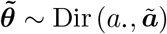, where 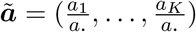.

Because of the relationship between the gamma and Dirichlet random variables, the propensity *θ_ijk_* can be represented as

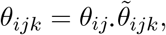

where

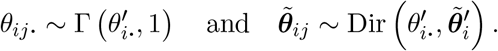

Likewise, we can represent 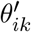 as

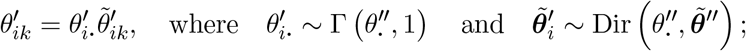

and we can represent 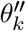 as

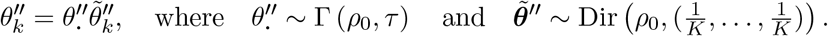

At each level in the hierarchy, we independently sample the sum from a gamma distribution and the proportions vector from a Dirichlet distribution The product of these yields a sample of the conditionally independent propensity value at that level.

This representation induces a hierarchical of Dirichlet prior over the proportions vectors at each level. Taking *K* → ∞, that hierarchical Dirichlet prior then describes the prior over the weights of the following hierarchical Dirichlet process (HDP):

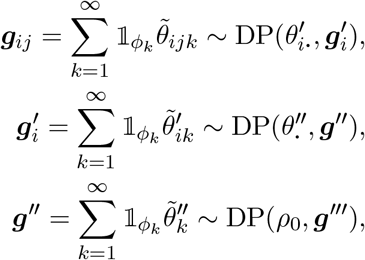

where *g*′″ is the base measure. This HDP prior is the same as the HDP prior in Section 2.2 except that the concentration parameters (e.g., *θ*″) are gamma random variables, as opposed to fixed hyperparameters, and are shared across all random variables at a given level.

## 4. Augment-and-Marginalize Gibbs Sampling for Gamma–Poisson Model

In this section the complete conditional distributions are derived for all latent variables in the Gamma-Poisson model—iteratively sampling from these constitutes a Markov chain whose stationary distribution is the exact posterior. The complete conditionals for all latent variables are available in closed form when further conditioned on a set of auxiliary variables. These auxiliary variables have closed form conditional distributions while leaving the stationary distribution of the Markov chain invariant; thus they facilitate efficient Gibbs sampling inference.

### 4.1. Latent subcounts

As with most Gamma-Poisson models, the first set of auxiliary variables that facilitate inference are the latent sub-counts *y*_*ijlb*1_,…, *y_ijlbK_* which sum to the observed count 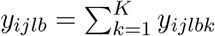. The *k*^th^ subcount *y_ijlbk_* represents the number of reads in sample *s* = (*i,j*) at locus *l* of base *b* that are allocated to latent subpopulation *k*. When conditioned on their sum, the vector of sub-counts is Multinomial-distributed:

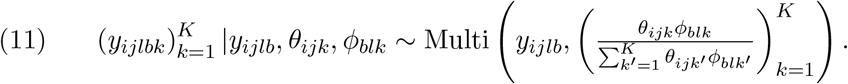

The complete conditionals of the other latent variables depend on different sums of these latent subcounts. Consider the total count of reads in sample *s* = (*i,j*) allocated to subpopulation *k*:

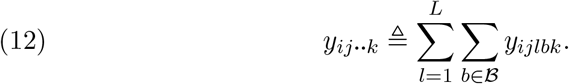

Due to the additive property of the Poisson distribution, this count is Poisson-distributed in the generative model:

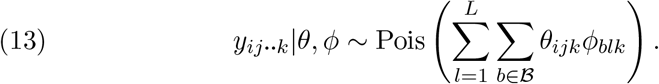

Since 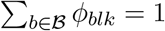 this simplifies to

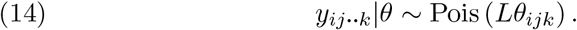

### 4.2. Augment-and-marginalize

Although this model posits a non-conjugate hierarchical Gamma prior, we can apply the “augment-and-conquer” procedure of Zhou and Carin (2012) to recursively marginalize out Gamma random variables and augment the model with auxiliary count variables to obtain closed-form conditionals for all latent variables. At a high level, the idea of augmentation is to represent a single complex distribution as a compound distribution such that when the compound distribution is appropriately marginalized the result is the original complex distribution. A simple example is the Student’s T distribution, which can be represented as a Gaussian distribution with an inverse Gamma prior on the variance parameter: when the variance is marginalized out, the result is a Student’s T distribution.

#### Marginalize θ_ijk_

The Poisson variable in Equation (14) represents the count of all reads whose distribution directly depends on *θ_ijk_*. Marginalizing out *θ_ijk_* gives a negative binomial distribution over *y_ij..k_*:

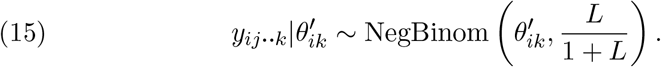

We note that previous work has shown that DNA sequencing count data is well-represented by a negative binomial distribution (Rabadan et al., 2018).

#### Augment with w_ijk_

If we now augment the model with the following Chinese Restaurant Table (CRT) random variable,

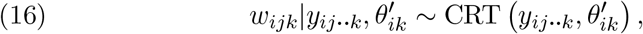

then the bivariate distribution 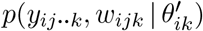 can be equivalently factorized as

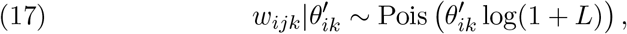

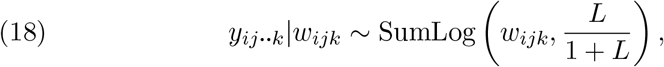

where SumLog (*w, p*) is the distribution of the sum of *w* i.i.d. Logarithmic random variables with probability parameter *p*. The Chinese restaurant table (CRT) distribution is the distribution of the number of nonempty tables in a Chinese restaurant process (Zhou and Carin, 2015). Suppose we have a Chinese restaurant process with concentration parameter *ρ*_0_ and *m* customers. Then, the number of occupied tables is 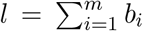 where *b_i_* ~ Bernoulli 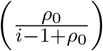 and the distribution of *l* is *l* ~ CRT (*m*, *ρ*_0_).

#### Inference in Augmented Model

A graphical model representation of the augment-and-marginalize procedure is shown in Figure 4. Figure 4a shows the original model structure with non-conjugate prior and Figure 4d shows the equivalent model structure where conjugacy holds. During inference, we sample the auxiliary variable *w_ijk_* using Equation (16). We may then proceed under the assumption that *w_ijk_* was in fact drawn from Equation (17) and that all dependence of 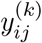 on 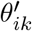 flows through *w_ijk_*. By marginalizing out *θ_ijk_* and augmenting with *w_ijk_* we have replaced a non-conjugate link from 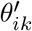 to *θ_ijk_* with a conjugate link from 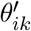 to *w_ijk_*. In the next steps, we recurse up the hierarchy.

**Fig 4:**
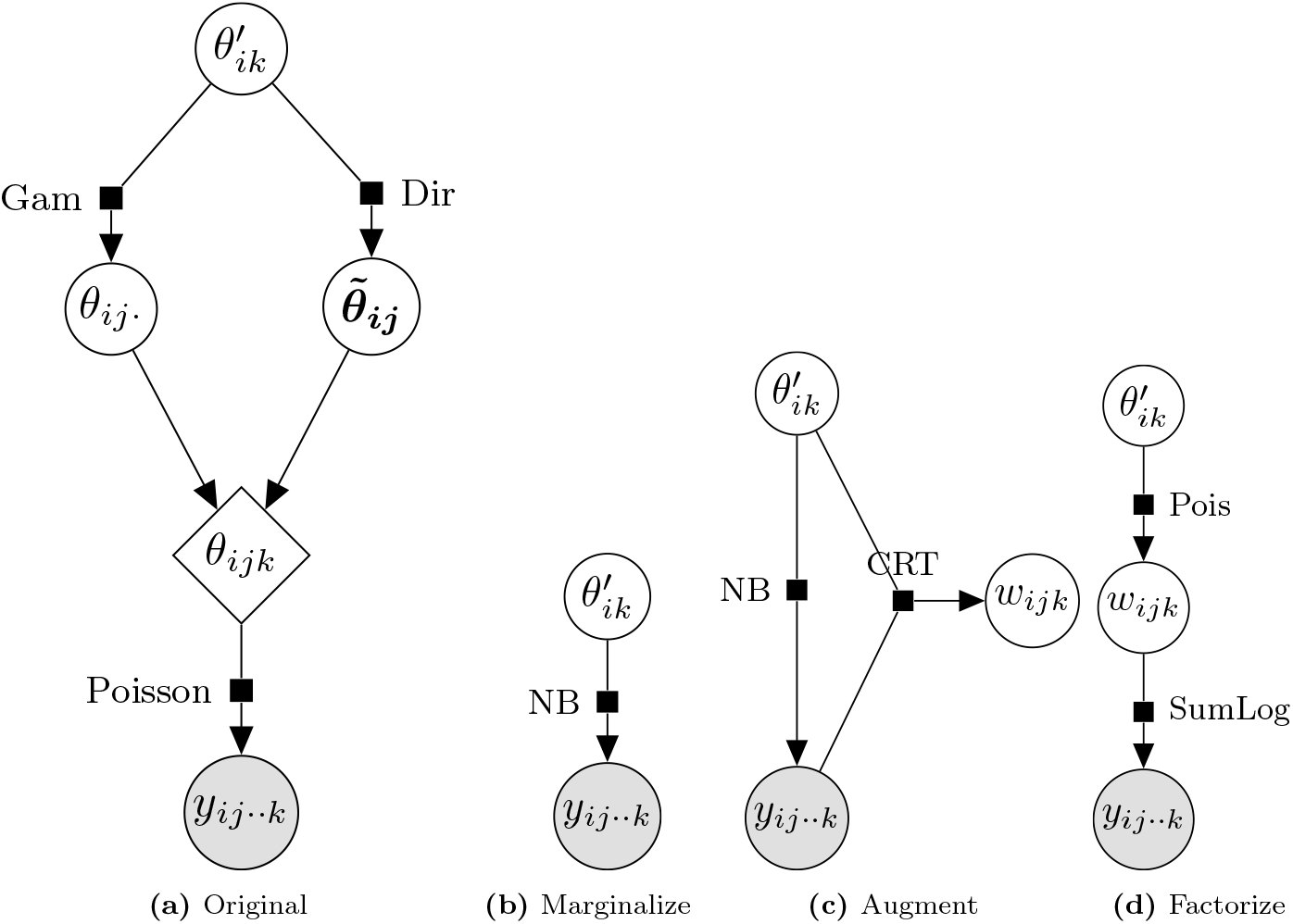
Augment-and-marginalize steps. (a) The original model structure. (b) The mod-elis transformed by marginalizing over *θ_ijk_*. (c) The model is augmented with the Chinese restaurant table (CRT) random variable *w_ijk_*. (d) Finally, the joint distribution 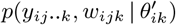 can be factorized using into the product of a Poisson and SumLog distribution.

#### Marginalize 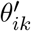

Having marginalized *θ_ijk_*, we now move up the model hierarchy to 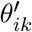. Define the 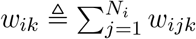 which is Poisson distributed,

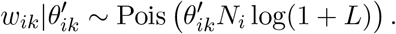

This count isolates the dependence of downstream variables on 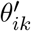, allowing us to marginalize 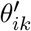 out—doing so induces the following negative binomial distribution over *w_ik_*:

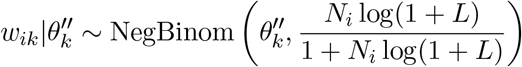

#### Augment with 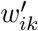

We augment the model with a CRT variable,

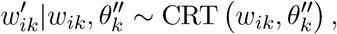

and then re-represent the bivariate distribution of *w_ik_* and 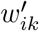 as:

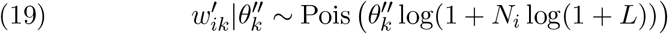

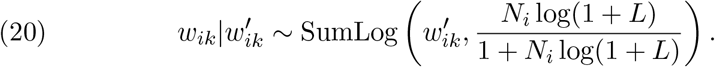

#### Marginalize 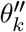

We now recurse up the hierarchy again. Defining the sum 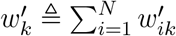, which is Poisson distributed:

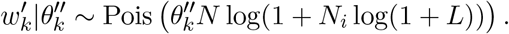

Marginalizing out 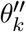 induces a negative binomial distribution:

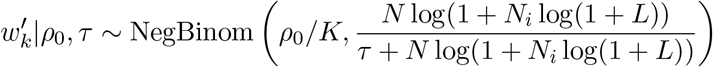

#### Augment with 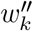

We augment the model with another CRT variable,

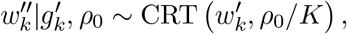

and then re-represent the bivariate distribution of 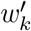 and 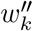 as:

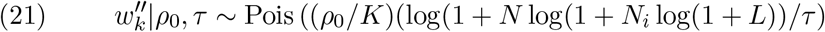

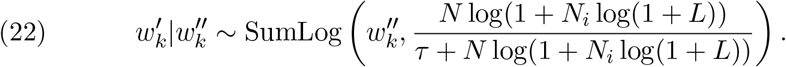

Doing so admits a conjugate link between *ρ*_0_ and 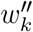.

### 4.3. Algorithm

The augment-and-marginalize derivation in the previous section involves introducing auxiliary variables which replace the nonconjugate links in the model with conjugate ones. This leads to an “upwards-downwards” Gibbs sampler in which we first sample auxiliary CRT counts up the hierarchy, then sample Gamma variables from conjugate conditionals down the hierarchy. A complete algorithm for the the Gibbs sampler for the Poisson-Gamma model is in Appendix C.

## 5. Simulation Experiments

In the previous sections we presented a novel hierarchical Dirichlet process mixture model for combining singlecell and bulk sequencing data and using the correlation between related samples induced by the sampling strategy or experimental design. The hierarchical Dirichlet process mixture model (Model hDP) allows for direct control over the a priori concentration of the sample, but the inference algorithm requires expensive Metropolis-Hastings steps. The hierarchical Gamma-Poisson model (Model hGP) can be interpreted as representation of the hierarchical Dirichlet process mixture model closely related to Model hDP with a much faster inference algorithm only requiring Gibbs sampling from analytical distributions. In this section, measure the accuracy, computational efficiency, and stability of these models compared to state-of-the-art machine learning and bioinformatics methods.

### Data Generation

Simulation data was generated from a parametric hierarchical Dirichlet mixture model with *K* = 3 true subpopulations and *L* = 5 genomic locations. The number of individuals is *N* = 6 and each individual has 1 bulk sample and 3 single-cell samples for a total of *N_i_* = 4 biopsies for each individual. The number of reads per sample (across 5 genomic locations) is *R_ij_* = 100; each genomic location has an average of 20 reads. Simulation data was generated according to the following model:

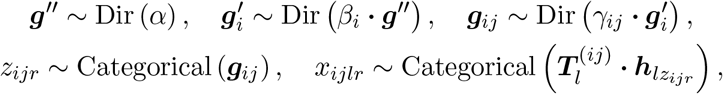

where *α* = (1,1,1), *β_i_* = 1, and *γ_ij_* = 10 for the bulk samples and *γ_ij_* = 0.1 for the single-cell samples. The genotype-nucleotide transition matrix is

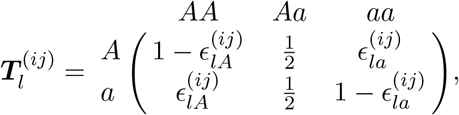

where 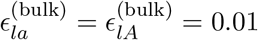 and 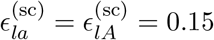 for bulk and single-cell samples respectively—(bulk) ≜ {*s* = (*i, j*) | *j* is a bulk sample} and (sc) ≜ = {*s* = (*i,j*) | *j* is a single-cell sample}. As a benchmark, the subpopulation-genotype matrix is set to

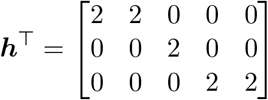

and in other simulation experiments ***h*** is randomly sampled from a prior distribution.

### Posterior Inference

The marginal posterior distributions ***g***″|***x, g***′|**x, g**|**x** and **h** were estimated using MCMC samples generated from Algorithm 1. We sampled 490 posterior samples after a burn-in/warm-up of 1,000 samples and thinning by a factor of 100. The number of subpopulations in the truncated Dirichlet process mixture inference algorithm (Algorithm 1) is set to *K* = 30 which is a factor of 10 greater than the true number of subpopulations. Figure 10 shows the true values and posterior distribution estimates from the simulation model, where the top three components of the model finds are exactly the same as the three components of **h** and the fourth component of the model finds has nearly zero probability. At population level, the KL divergence between the true distribution and inferred posterior distribution is 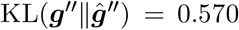. At individual level, the average KL divergence is 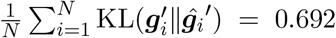 and the standard deviation is 0.289. At the biopsy level, the average KL divergence is 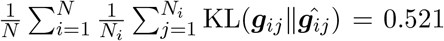 and the standard deviation is 0.371. A detailed visualization and discussion of the posterior distribution are in Appendix D.1. These results indicate that the model is able to identify the true subpopulation genotypes and the posterior distributions are appropriately uncertain relative to the amount of data and the proximity to the data in the model.

### Comparison to LDA and NNMF

To assess the importance of the hierarchical structure in Model hDP, we compared the performance to latent Dirichlet allocation (LDA) and non-negative matrix factorization (NNMF). However, both LDA and NNMF failed to find the true components for our benchmark simulation data. Thus we generated other data sets using the same parametric model but only having bulk data and compared the performance with our model. The KL divergence, 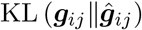, mean and 95% confidence interval across inference repeats is shown in Figure 11 in Appendix D.2 (Table 2 in Appendix D.2 shows the numerical values). These experiments shows that Model hDP outperforms LDA and NNMF for bulk-only and mixed data scenarios.

**Table 1.**
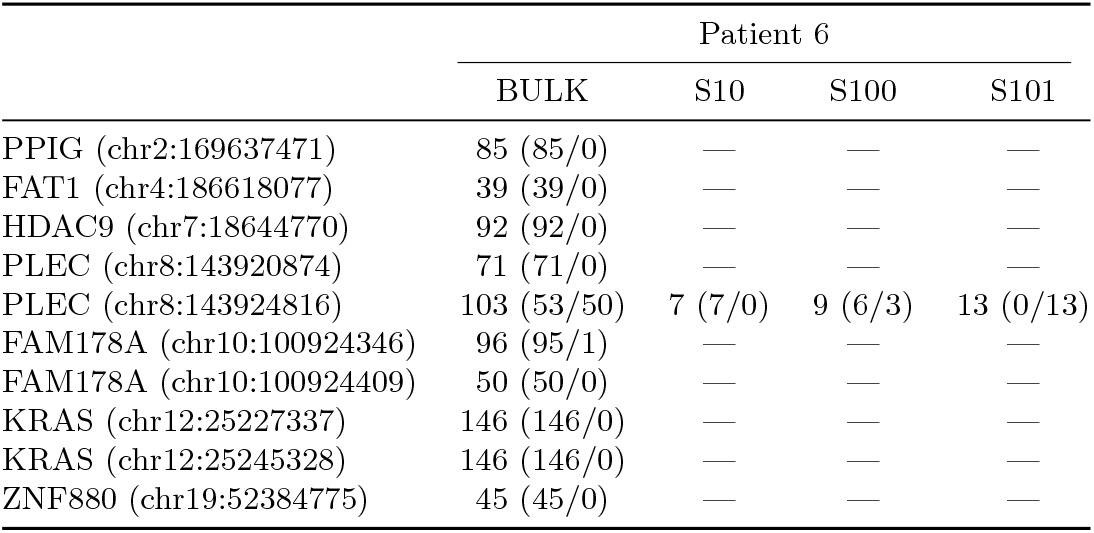
Read-count table for Patient 6. The total read counts across ten loci for one bulk sample and three single-cell samples (S10, S100, S101) are shown. The major/minor allele ratios are shown in parenthesis after each read count. Zero read counts are shown as dashes indicating missing data at those loci.

**Table 2:** Comparison between LDA, NNMF, and Model hDP by KL divergence between estimated sample-level posterior distribution over components and true distribution over components, 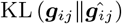.

| N | N<sub>i</sub> | LDA | NNMF | hDp Model |
| --- | --- | --- | --- | --- |
| 3 | 3 | — | 0.035 (0.035, 0.036) | **0.009 (0.004, 0.013)** |
|  | 10 | 0.208 (0.208, 0.208) | 0.035 (0.033, 0.037) | **0.011 (0.009, 0.013)** |
| 10 | 3 | 0.235 (0.235, 0.235) | **0.002 (0.001, 0.002)** | 0.016 (0.000, 0.036) |
|  | 10 | 0.276 (0.276, 0.276) | 0.012 (0.012, 0.012) | **0.008 (0.004, 0.012)** |
| 20 | 3 | 0.198 (0.198, 0.198) | 0.019 (0.018, 0.020) | **0.007 (0.006, 0.008)** |

### Comparison to PhyloWGS and TreeClone

We compared the performance of Model hDP to two other models for inferring clonal subpopulations in mixed samples: PhyloWGS and TreeClone. Due to sample size limitations in PhyloWGS, we reduced the sample size to *N* ∈ {1, 2}), *N_i_* ∈ {1, 2, 3} and only generated bulk data and kept other settings same (*R_ij_*=100, *L*=5) to generate 6 simulation datasets to compare PhyloWGS, TreeClone and Model hDP. PhyloWGS successfully identifies true subpopulations at sample level but inferred an incorrect hierarchical structure relating the samples. A complete presentation of the results of this experiment are in Appendix D.3. Figure 12 therein shows phyloWGS’s posterior distributions at sample level are less accurate than Model hDP in terms of KL divergence. TreeClone successfully identifies number of subpopulations but the posterior distributions at sample level are less accurate than our model.

### Sensitivity Analysis

To assess the sensitivity of the model, we performed simulation experiments varying ***h**, K, L*, and the single-cell sequencing error rate 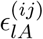 and 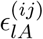. To assess the sensitivity to different values of ***h***, five simulation data sets were randomly generated with ***h**_lk_* ~ Multi(1, ***a***) where ***a*** = (0.45, 0.1, 0.45). In all cases, the model was able to identify the true subpopulation genotypes when the subpopulation was represented by samples in the data as shown in Appendix E.1. To assess the sensitivity to varying *K, K* was increased from 3 to 10 with L = 10 and *R_ij_* = 1000 with randomly sampled ***h***. Appendix E.2 shows the results of the simulation experiment of vary *K*: our model successfully inferred true components with relatively small KL divergence. To assess the sensitivity to varying *L*, we ran simulation experiments with *L* = {3, 10, 20, 50, 100}. Appendix E.3 shows our model identified the subpopulations and posterior distribution of the subpopulations across this range of *L*. Finally, we assessed the sensitivity of the model to varying single-cell sequencing error rate 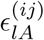 for the single-cell samples because while the error rate for bulk experiments is well-established, the error rate for single-cell experiments is more uncertain. We set 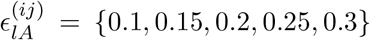 and sampled ***h*** randomly. The results (Appendix E.4) show our model was not sensitive to the specified singlecell sequencing error rate, but performance does degrade as the error rate increases.

### Accuracy and Computational Efficiency of Model hGP

Since Model hGP is designed for a large number of single-cell samples, we simulated *N_i_* = 99 single-cell samples from the benchmark model with a sequencing error rate of 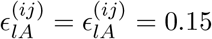. The model took less than 10 minutes to run and identified all three subpopulation genotypes exactly, 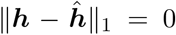. The accuracy of marginal subpopulation distributions were, on average: 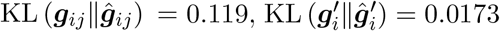, and 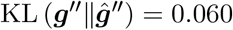. These metrics indicate that Model hGP is highly accurate and computationally efficient for the intended use regime—a dataset with a large number of single-cell samples.

## 6. Acute Lymphoblastic Leukemia Experiments

We fit Model hDP and Model hGP to a mixed single-cell and bulk DNA sequencing data set of *N* = 6 childhod acute lymphoblastic leukemia (ALL) patients (Gawad, Koh and Quake, 2014). The study collected *targeted sequencing* of a panel of SNV loci from 1,479 single-cells and bulk samples to better understand genomic heterogeneity. The authors of that study concluded that *KRAS* mutations occur late in development, but do not lead to clonal takeover. DNA sequencing data from bulk samples and single-cell samples was obtained from the NCBI short read archive under study accession SRP044380.

### 6.1. Preprocessing

Sequenced reads from both bulk and single cells were converted to FASTQ format and mapped to the human genome assembly (hg38) using the Burrows-Wheeler Alignment tool (BWA version 0.7.17) with default parameters to create BAM files (Li and Durbin, 2010). The reads with mapping quality below 40 were removed and PCR duplicate marking was performed with Picard (version 2.0.1). The results of the preprocessing can be tabulated as shown in Table 1 for Patient 6 for one bulk sample and three single-cell samples. It is evident that there is good coverage across all of the loci for the bulk sample, but the single-cell coverage is both sparse and shallow. Other patient samples (shown in Appendix F.4) have single-cell coverage at different loci. The models we have developed address this single-cell sparsity issue by borrowing strength across patients, bulk samples, and single-cell samples to provide a more accurate picture of the genetic state of the samples, patient, and population.

#### 6.2. Posterior Inference using Model hDP

Model hDP is most appropriate for for targeted sequencing experiments (small *L* and small 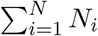) and a mixture of bulk and single-cell experiments because it allows one to add impactful a priori information about the data in the hyperparameters of the model when the sample size is small. We sampled three single cells and one bulk sample for each patient. Of the mutations validated in the original report, we selected *L* = 10 non-synonymous loci curated from ALL literature for which there was read support in both the bulk sample and at least one single cell. This setup replicates a scenario where one has a biomarker panel for targeted therapeutic decision-making while employing the published data set. A full listing of the loci and samples selected for analysis is given in Appendix F.1 and Appendix F.2. We set *K* (the number of subpopulations in the model) to 30 which we expect is much greater than the number of true subpopulations based on literature on genomic subtypes. We set the parameters *α* to 1, *β_i_* to 1, and *γ_ij_* to 0.1 for all *i* and *j*. Single-cell samples are amplified by whole-genome amplification the nucleotide error rates are expected to be much higher than for bulk samples (Zafar et al., 2016), so the different error rate models are employed for bulk and single-cellsamples at all locus positions *l* = 1, …, *L*,

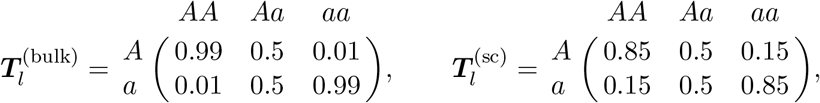

where (bulk) = {*s* = (*i, j*) | *j* is a bulk sample} and (sc) = {*s* = (*i, j*) | *j* is a single-cell sample}. We drew a total of 50,000 samples with a burn-in of 1,000 samples and thinned by a factor of 100 giving 490 posterior samples. Convergence of the sampler was validated by standard Geweke tests (see Appendix F.3). With these parameters, inference took 7 hours on a single processor core. In further testing, we found that in fact 50,000 posterior samples are not required and 10,000 samples would achieve similar results, thus the time can be reduced by a factor of five. Since ***h*** is discrete, at the end of the sampling process we align samples of ***h*** scan across all of the samples to register unique ***h**_k_* with associated components of 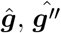, and 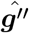. For example, suppose subpopulation 3 in MCMC sample 100 is 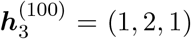, and in MCMC sample 143 subpopulation 6 is 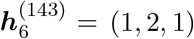. Clearly, the subpopulations in both samples are the same, so we associate 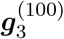 with 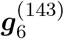.

#### Posterior Distribution

The posterior distribution as estimated from the samples is concentrated on only a few subpopulations indicating that the truncated Dirichlet process used for the inference algorithm is an accurate approximation. Figure 6 shows the average of the all 490 samples of posterior distribution over populations, subpopulations and bulk and three single-cell samples of Patient 6. We selected subpopulations with an average posterior greater than 0.05 in any sample from Patient 6, 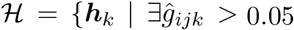, for *i* = 6 and *j* = 1, 2, 3, 4}. The y-axis is the MCMC estimate of 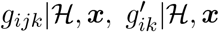, and 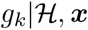. Similar plots for all six patients can be found in Appendix F.4.

**Fig 5:**
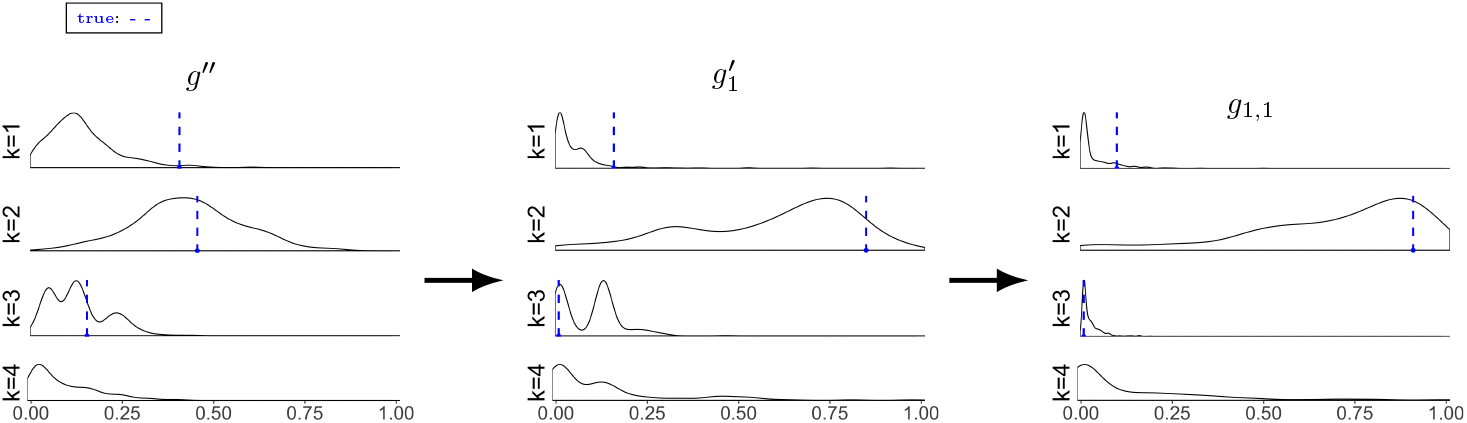
Marginal posterior density estimates for simulation data with *N* = 6 and *N_i_* =4. The population-level subpopulation marginal distribution ***g***′ slightly underestimates the fraction of subpopulation *k* = 1 and is accurate for the other subpopulations. The accuracy of the individual-level distribution 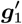 and the sample-level distribution ***g***_1,1_ are more accurate in part because because they are closer to the data in the model hierarchy.

**Fig 6:**
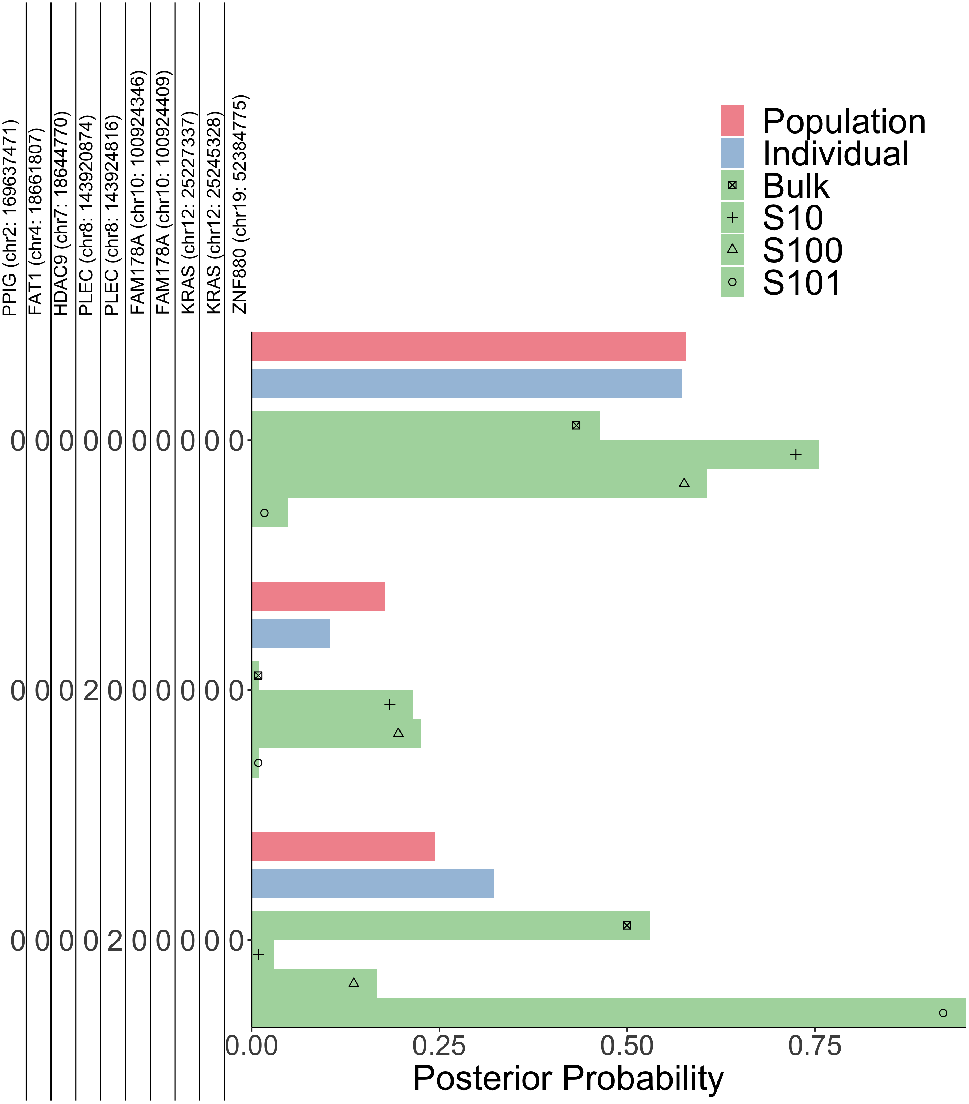
Posterior distribution plots for Patient 6 from Model hDP. Red bars show the population level distribution over subpopulations 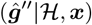, blue bars show the individual level distribution 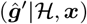, and green bars show the sample level distributions 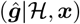, where 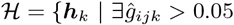, for *i* = 6 and *j* = 1, 2, 3,4}.

The posterior distribution is smoother at the population level than at the sample level reflecting the smoothing effect of the hierarchical model structure. At sample level, most of the posterior distributions are concentrated at one component. As can be seen in the figures in Appendix F.4 bulk samples tend to be more mixed than single-cell samples reflecting the biological reality that single-cell samples contain only one genotype, while bulk samples are a mass of cells each with their own genotype.

A useful feature of the hierarchical model is its ability to share information across samples through the individual and its ability to share information across individuals through the population. This feature is particularly powerful for single-cell data where the read coverage may be zero for some loci because the model can rely on the individual-level distribution which is informed by both the population distribution and the bulk sample. Table 1 shows a read-count table for Patient 6. While all of the loci have data from the bulk sample, only one locus, *PLEC* (chr8:143924816), has any single-cell data. This table is representative of single-cell and bulk data in that bulk samples tends to have good coverage across all loci, while single-cell samples tends to have much more missing data (see Appendix F.4). Recalling the posterior distribution in Figure 6, the bulk, S10, and S100 samples all have significant posterior mass on the wild-type (all-zero) component, but S101 has very little mass on that component which is consistent with the read-count data in Table 1. The posterior distribution places some mass on components with a homozygous mutation in *PLEC* (chr8:143920874) and *KRAS* (chr12:25227337) for single-cell samples S10 and S100. Of course, single-cell samples are expected to have the posterior mass concentrated on only one genotype. Table 1 shows that there is missing data for these loci and because the data is missing the model is employing information from the individual-level distribution which places roughly similar mass on those components. This model behavior is consistent with our expectation that a lack of data is not evidence of no mutation, but instead should be informed by the information from the bulk through the individual-level distribution.

#### Biological Interpretation

As shown in Figure 6, for Patient 6, the posterior distributions of the bulk data and single cell S101 have more than 50% probability on the subpopulation which has a single mutation at *PLEC* (chr8: 143924816) at the sample level. The posterior distribution for single cell S101 has more than 75% on that component. This result correlates with the description of the data in the original report (Gawad, Koh and Quake, 2016). Model hDP also finds meaningful mutations for Patient 1-5 after comparing our result with the original report. As shown in Appendix F.4, Patient 2 has a posterior distribution that concentrates on the component that has a mutation on *PLEC* at sample level which is exactly the same as described in the original report. For Patient 5, the posterior distribution at sample level for S10 has nearly 50% probability concentrated on two components both with a mutation on *FAM178A.* This indicates a possible mutation on *FAM178A* for Patient 5 which is congruent with the original paper. The posterior distribution for S100 has a large probability concentrated on the component with a mutation on *HDAC9.* It is expected since all two reads on *HDAC9* harbor a minor allele. For Patient 4, the posterior distribution at sample level is concentrated on the component that has a mutation in *FAT1* which is shown to be a gene that has mutated in pediatric ALL patients (Neumann et al., 2014). Our model also finds an interesting mutation in *PPIG* in Patient 1; it is not yet known if the mutation acts to drive ALL development, but the mutation is clearly present in a single cell.

### 6.3. Posterior Inference using Model hGP

The hierarchical GammaPoisson model (Model hGP), has a fast inference algorithm and therefore, can be used to analyze the ALL data set. We selected all the single-cell data for this analysis giving 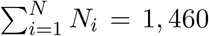 samples and *L* = 111 non-synonymous loci from 6 patients. *SERPINF2, RNF180* are found mutated across all patients remove form the analysis giving *L* = 109 loci. Inference on this data set with Model hGP took 80 minutes on a 4-core MacBook Pro with 16GB RAM and 2.3GHz processor using a Cython implementation.

#### Posterior Distribution

Figure 7 shows the posterior probability—under population-level, individual-level, and sample-level distributions—of the five subpopulations that had the highest posterior probability under Patient 1’s sample-level distributions and three single cells. It is evident there is heterogeneity in the clonal content of the tumor. Single-cell 1 has a large posterior weight on subpopulation 1, single-cell 2 has a large weight on subpopulation 2 and single-cell 3 has a large weight on subpopulation 4. Each subpopulation is associated with a genotype given by ***ĥ***. Figure 25 in Appendix F shows the ***ĥ*** matrix for the subpopulations in Figure 7.

**Fig 7:**
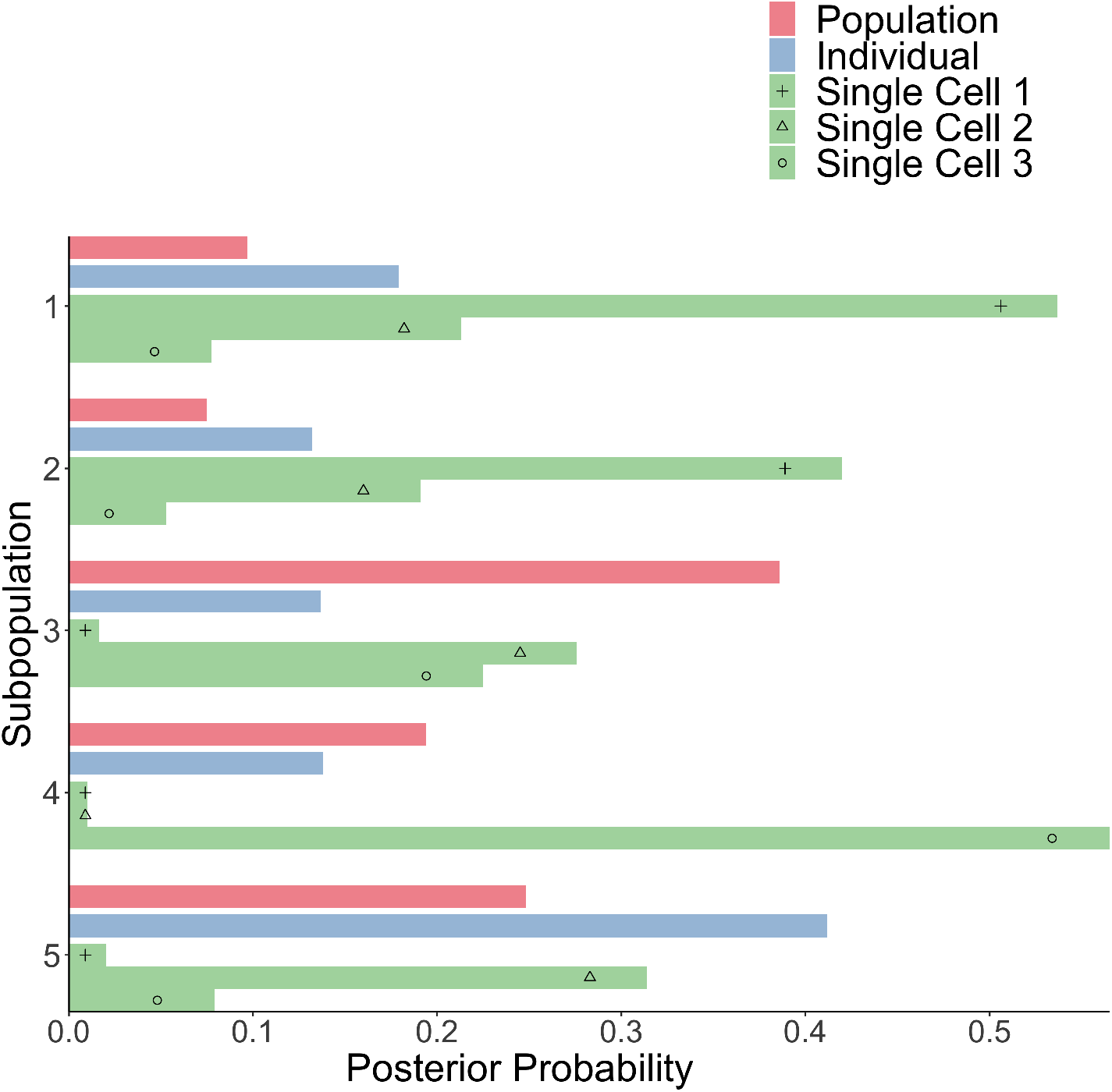
Posterior distribution plots for Patient 1 for Model hGP. Red bars show the population level distribution over subpopulations 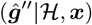, blue bars show the individual level distribution 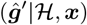, and green bars show the sample level distributions 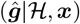.

#### Biological Interpretation

One way Model hGP can be used to draw inferences that are not obvious from direct inspection of the data is to infer the co-occurence of mutations across samples. If two genes are frequently mutated together it may indicate a synergistic relationship between two oncogenic processes mediated by the genes. An *L × L* adjacency matrix, ***A***, can be constructed from the model as where an element is

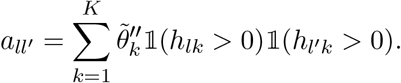

The adjacency matrix values are bounded between zero and one and a large value indicates that the two loci are co-mutated and have a high posterior probability across samples.

Figure 8 shows the adjacency matrix in network form where an edge between *l* and *l*′ is drawn if *a_ll′_* > 0.50. Loci without edges to other loci are omitted. There are 67 mutations that meet the criteria for inclusion. The most connected locus has 18 connected loci and the average number of connections is 5.

**Fig 8:**
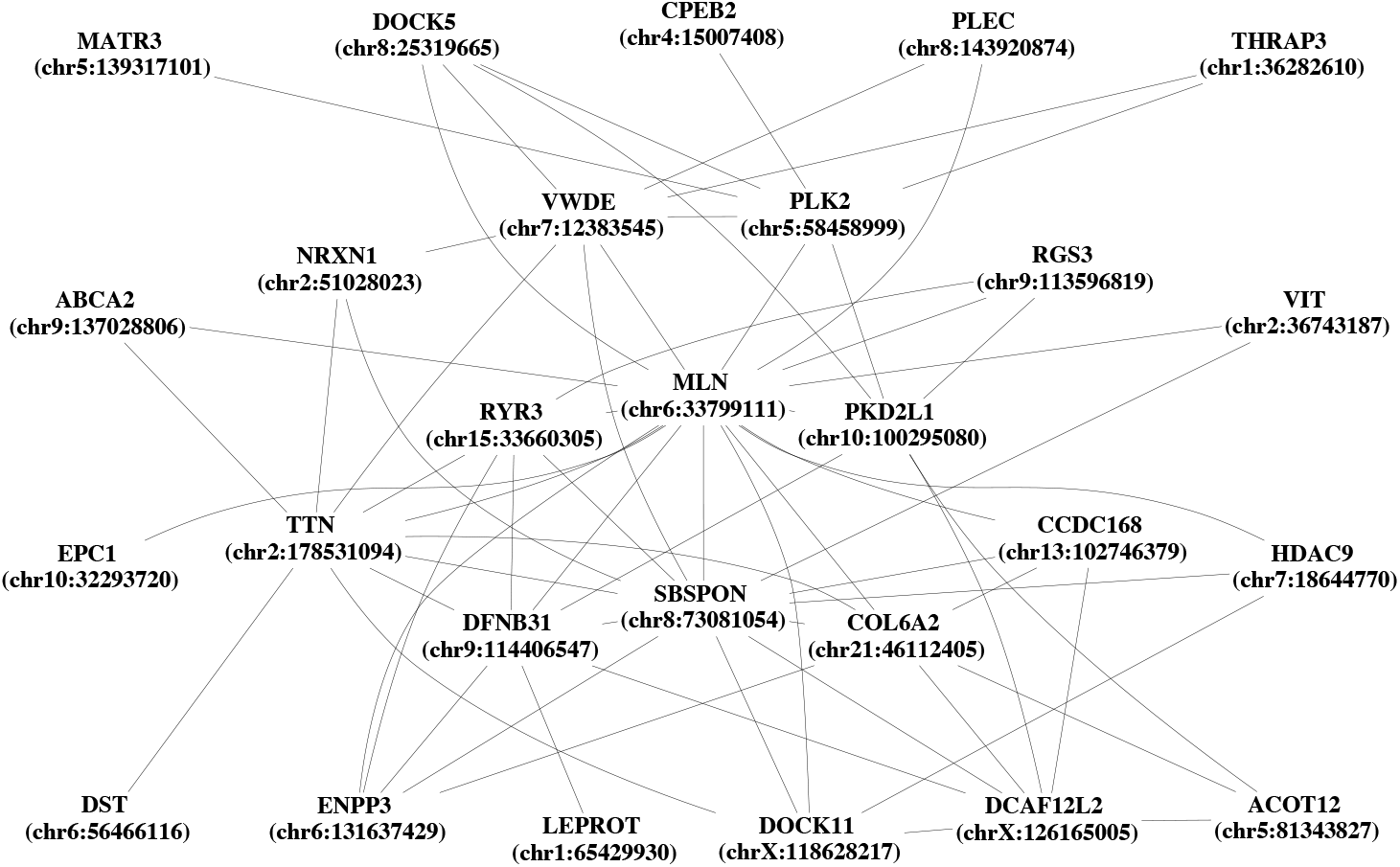
Inferred mutation co-occurence network across all patients from Model hGP.

The most connected component is *MLN* (chr6:33799111) and all of the reads associated with mutations in *MLN* occur in single-cell samples from Patient 5. One of the highly connected genes, *TTN,* has recently been reported as the most frequently mutated gene in a pan-cancer cohort and is associated with increased tumor mutation burden (Oh et al., 2020). Interestingly, *TTN* is connected to *RYR3* and this connection was identified in the original report (Gawad, Koh and Quake, 2016) in the inferred directed minimum spanning tree of subpopulation evolution for Patient 1. *TTN* was identified as the founder mutation in Patient 3 and a downstream mutation *DST* is also shown to be connected in our inferred network. Though it should be noted that *TTN* is a very large 304kb gene. *PLK2* is connected to seven other loci including *DOCK5.* This co-occurence was also observed in the original report for Patient 4 (Gawad, Koh and Quake, 2016).

The co-occurence adjacency matrix, **A**, can be constructed with only data from an individual (patient). Figure 9 shows the adjacency matrix for Patient 1. In this network *MLN* (chr6:33799111), *TTN* (chr2:178531094), *VWDE* (chr7:12383545) and *PLK2* (chr5:58458999) are the most connected mutations. While these co-occurence inferences are suggestive, and not conclusive they are powerful for proposing avenues of validation through observational human data or experimental model systems.

**Fig 9:**
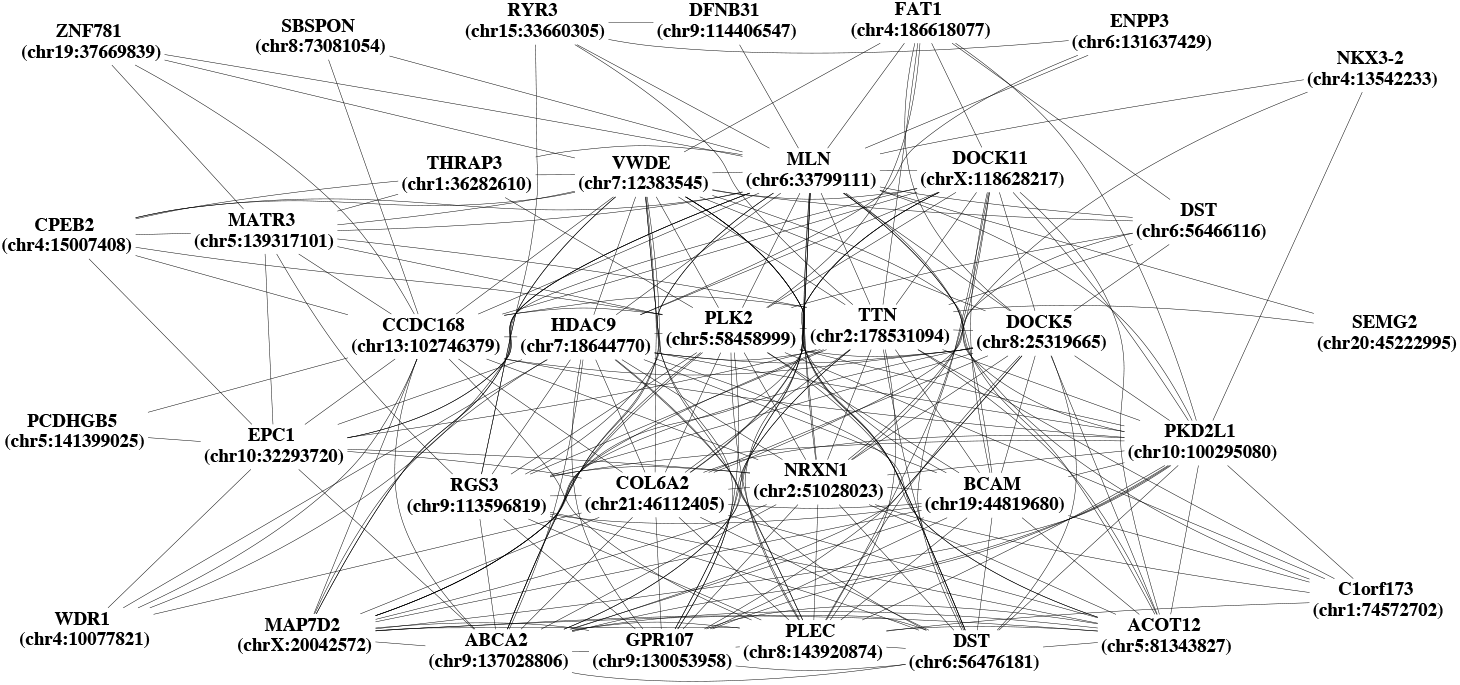
Inferred mutation co-occurence network across Patient 1 from Model hGP.

## 7. Discussion

We have presented a novel statistical model and companion inference algorithms for inference in structured single-cell and bulk DNA sequencing data. We have also suggested an alternative representation of the model as a Gamma-Poisson hierarchical model. The reason for the development of two inference algorithms is that they each make different tradeoffs between computational efficiency and statistical representation. The Dirichlet process model has parameters *γ_ij_* and 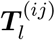 that can be individually specified for each sample *s* = (*i,j*). A single-cell sample would have a small value of *γ_ij_* indicating there is, a-priori, a small number of subpopulations and the genotype-nucleobase transition matrix, 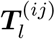, may be set according to an error model of DNA sequencing data after whole genome amplification. A bulk sample, conversely, may have a larger value of *γ_ij_* and a value of 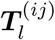 with a lower sequencing error rate. This representational flexibility in the model comes with computational costs and the MCMC inference procedure can be slow for a MCMC sampling algorithm are more relevant for targeted sequencing experiments on small study populations—for example correlative sequencing data in a phase I clinical trial. The Gamma-Poisson model does not provide direct control over the concentration of subpopulations for individual samples and the parameter 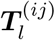 is the same for all samples. This limitation may not be critical when sufficient data exists to achieve accurate inferential results from the entire data set or when the experimental protocol is the same for all samples. Inference Gamma-Poisson model uses analytical updates in a Gibbs sampler and is very fast making it feasible to analyze larger data sets. The Gamma-Poisson model and augment-and-marginalize Gibbs sampling algorithm are more relevant for sequencing experiments on large study populations with single-cell samples.

The inference algorithm for the Dirichlet process mixture model is highly accurate compared to standard decomposition models and existing bioinformatics tools for structured, targeted sequencing data sets. An analysis of a real sequencing data set reveals the inferred genotypic content of the sample and the a-posteriori distribution over clonal subpopulations with associated uncertainty based on incomplete single-cell sequencing.

An analysis of a large-scale sequencing experiment using this model revealed co-occurence networks for each individual patient. Some co-occurence connections were hinted at in the original report of the data set, confirming the ability of the model to identify connections in the co-occurence network. The Gamma-Poisson model provides a more comprehensive and unbiased analysis of that data set by combining evidence from all of the data under a Bayesian nonparametric hierarchical model.

Copy number aberration is a prevalent in cancer samples and an important aspect of cancer etiology and statistical inference in genomic data. We have assumed in this work that the samples are diploid with no copy number aberration. There are technologies to independently measure copy number aberration (Alkan, Coe and Eichler, 2011) and methods for estimating copy number aberration from sequencing data (Budczies et al., 2016). Some methods have demonstrated ability to jointly estimate single-nucleotide variants and copy number aberrations (Riester et al., 2016) and such joint estimation would be interesting future work for the models developed here.

## Acknowledgements

We would like to thank Alexandre Bouchard-Côté and Mingyuan Zhou for reading an early draft of this paper. This work was supported by NIH 1R01GM13593101.

## APPENDIX A: RELATED WORK

### Paired Tumor-Normal Models

There has been substantial work on analyzing tumor heterogeneity using paired tumor-normal samples. PurityEst (Su et al., 2012) and PurBayes (Larson and Fridley, 2013) focus on estimating the purity of a sample, but fundamentally assume the mixture in a sample is exclusively between a tumor genotype and a normal (non-tumor) genotype. *Pyclone.* Roth et al. (2014) proposed a Dirichlet process mixture model for subpopulations called Pyclone. Pyclone is inspired by phylogenetic considerations, but the actual subconal populations are not constrained to agree with a tree. The method has enjoyed considerable success in applications (Andor et al., 2016; McGranahan et al., 2016).

### PhyloWGS

PhyloWGS uses a Bayesian nonparametric model to reconstruct genotypes of the subpopulations from sequencing data (Deshwar et al., 2015). This was one of the first models to attempt to reconstruct both the point mutation landscape as well as the copy-number variation landscape for complex tumors. The paper shows that copy-number variation data is essential for accurate subclonal reconstruction. However, that work did not look at the effect of incorporating the experimental design structure or single-cell sequencing.

### Bayclone

Bayclone uses an Indian buffet process prior over the genotypes for the subpopulations (Sengupta et al., 2015). While Bayclone focuses on the subpopulation genotypes, it also incorporates a Dirichlet distribution for the subpopulation fractions in each sample. However, it assumes each sample has the same probability for each non-normal subpopulation a-priori, and it assumes each sample is conditionally independent given this prior. Bayclone is related to the phylogenetic Indian buffet process—a feature allocation model (Miller, Griffiths and Jordan, 2008).

### Sciclone

Sciclone uses a hierarchical Bayesian mixture model to infer sub-clonal populations (Miller et al., 2014). The method achieves computational efficiency by using a variational approximation to estimate the model parameters. It uses a pruning method to select the number of subpopulations and provides an estimate of uncertainty in the inferential products.

### CloneHD

CloneHD integrates information from copy number data, B-allele frequency, and somatic nucleotide variants to infer clonal subpou-lations (Fischer et al., 2014). The method uses the Bayesian information criterion to select the number of subclonal populations. It uses a coupled hidden Markov model to integrate data across omic modalities.

### Clomial

Clomial proposes a hierarchical model that has Bayesian conjugacy and therefore closed form EM updates (Zare et al., 2014). Their experiments show that even without information about the proximity of sampling within a tumor, nearby samples display similar clonal composition as would be biologically expected.

### Cloe

Cloe takes the innovative approach of incorporating a prior over phylogenetic trees (Marass et al., 2016). While the prior regularizes the resulting products of inference towards valid phylogenetic tree structures. The model requires a somewhat costly Metropolis-coupled Markov-chain Monte Carlo sampler, but for targeted sequencing, the expense is not prohibitive.

### TreeClone

Treeclone is a nonparametric Bayesian model for reconstructing the clonal subpopulation phylogeny and inferring tumor heterogeneity (Zhou et al., 2019a). It employs a tree-based latent feature allocation model on pairs of mutations (Zhou et al., 2019b) that are phased by their presence on the same short-read. By constraining the columns of the mutation-pair-by-subclone matrix to a tree structure MCMC sampling is much more computationally efficient. The method produces impressive results for a moderate number of samples and scales well with the number of mutation pairs.

## APPENDIX B: HIERARCHICAL DIRICHLET PROCESS MIXTURE MODEL MCMC SAMPLER DERIVATION

Derivations for each sampling step in the MCMC sampler for the Model hDP are provided in this section. The index of the MCMC sample is denote by a superscripted (*t*).

### Sample 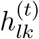 from p(h_lk_|**h**_-lk_, **x, z, a**_l_, **T**)

The matrix **h** can be sampled by updating its elements independently conditional on the Markov blanket. The posterior distribution of *h_lk_* is

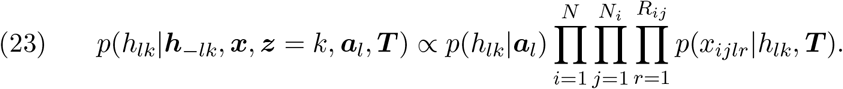

The conditional term *z* = *k* denotes the set of reads assigned to subpopulation *k*, {(*i, j, r*)|*z_ijr_* = *k*} since *h_lk_* only depends on the reads assigned to subpopulation k. The normalization constant can be computed using the constraint 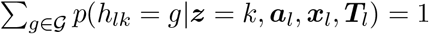. A sample *h_lk_* is drawn from a multinomial (categorical) distribution.

### Sample 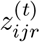 from p(z_ijr_|z__ijr_, **x, h, g**)

The matrix ***z*** can be sampled by sampling each *z_ijr_* due to the conditional independence structure of the model. The posterior distribution of *z_ijr_* is

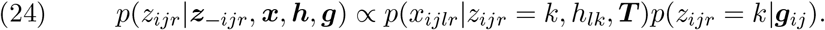

The terms *p*(*x_ijlr_*|*z_ijr_* = *k*, *h_lk_*, **T**)*p*(*z_ijr_* = *k*|***g**_ij,j_*) can be computed exactly for each *k* = 1,…, *K*. These quantities are normalized to give *p*(*z_ijr_*|***z**_-ijr_*, **x, h, g**). A sample *z_ijr_* is drawn a categorical distribution with associated probabilities.

### Sample 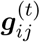 from *p*(***g**_ij_*|***g**_-ij_*, **z, g**′,*γ_ij_*)

The sample-level distributions over subpopulations, ***g***, can be sampled by updating each ***g**_ij_* because the ***g**_ij_*’s are conditionally independent given 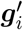. The posterior distribution of ***g**_ij_* is

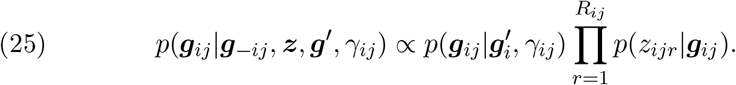

The likelihood is a categorical distribution with *K* possible components and the prior is a *K* dimensional Dirichlet distribution; by Bayesian conjugacy, the posterior distribution is the *K* dimensional Dirichlet distribution:

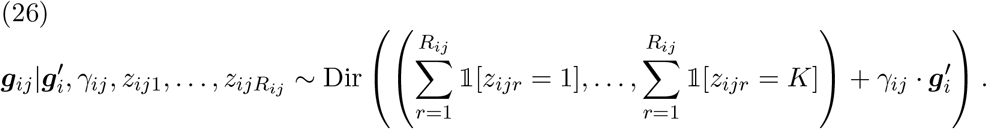

### Sample 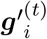 from 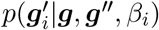

The set of distributions of subpopulations for all individuals ***g***′ can be sampled by sampling each 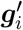 independently conditioned on 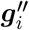. The posterior distribution of 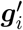 is

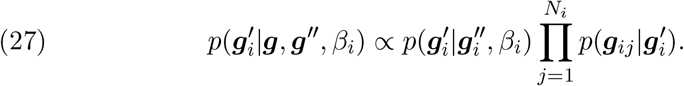

We use a simple Metropolis-Hasting sampler to draw from the posterior distribution because the prior and likelihood are not Bayesian conjugates.

### Sample **g**″ from p(**g**″|**g**′, **g**′″, α_0_)

The posterior distribution of subpopulations in the population (entire dataset), ***g***″, is

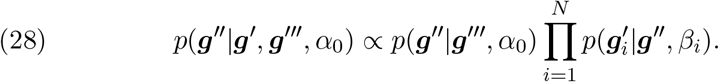

The prior is 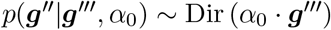 and the likelihood is 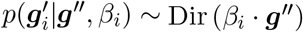. So, we use a Metroplis-Hasting sampler to draw a new sample ***g***″.

## APPENDIX C: GAMMA-POISSON MODEL AND INFERENCE

A complete derivation of the Gibbs sampling steps and associated notation for the Gamma-Poisson model is shown in the main text. Here we summarize that algorithm in Algorithm 2.

**Algorithm 2:**
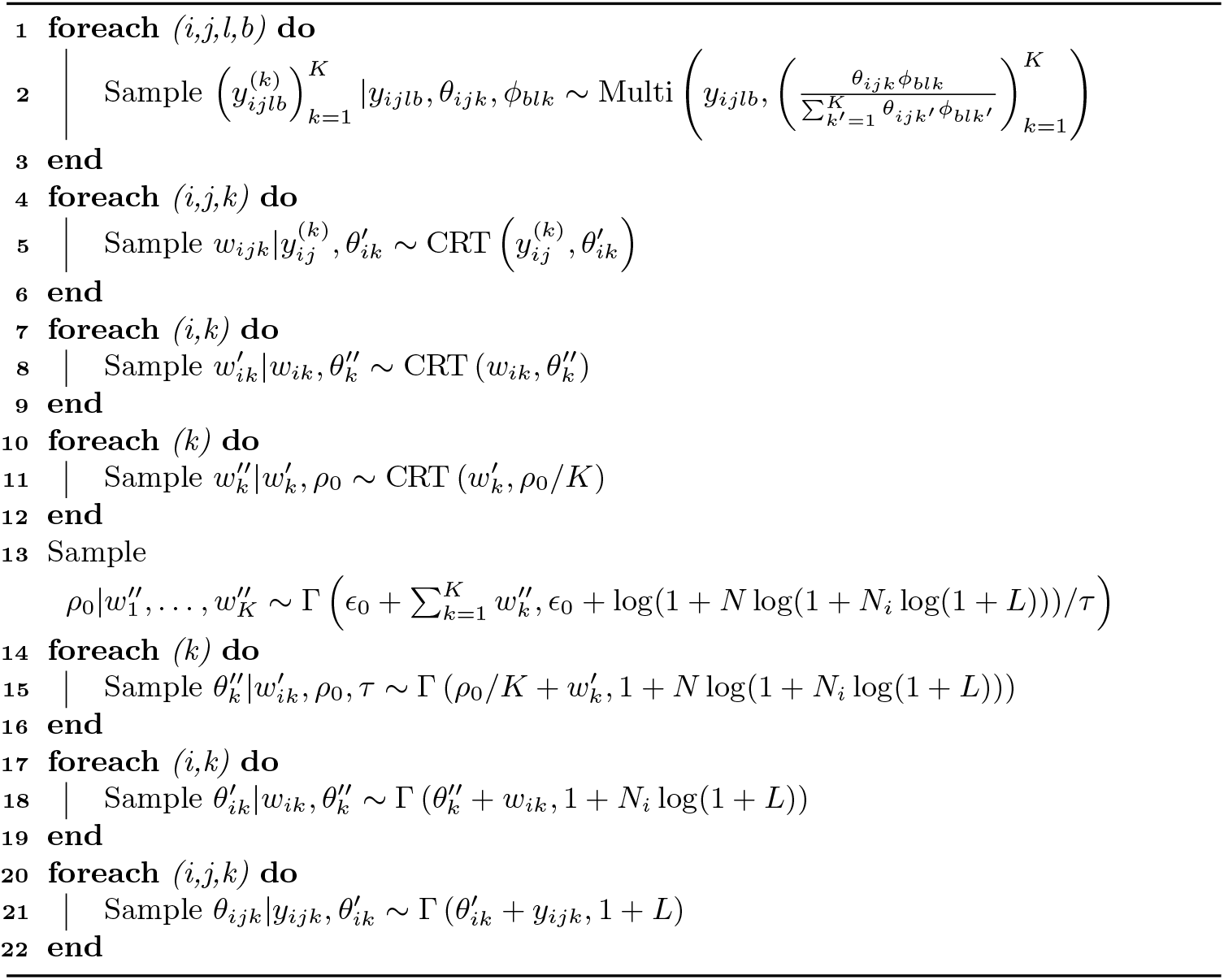
Auxiliary variable Gibbs sampler for Gamma-Poisson Model

## APPENDIX D: SIMULATION EXPERIMENTS

### D.1. Posterior Inference

As described in the main article, the model identifies the true number of subpopulations by placing most of the posterior mass only on three subpopulations. The fourth most frequent subpopulation (*k* = 4) is shown as evidence that the subpopulation is not employed by the model. At the sample level (***g**_ij_*) the posterior inference is highly accurate for both pure and mixed samples. At the individual level 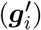, the posterior distribution displays more uncertainty (less peaked), though the posterior mode is still accurate. At the population level (***g***″), the posterior distribution is more uncertain because it is furthest from the data in the hierarchy and closest to the prior but still reasonably accurate considering the small number of individuals (*N* = 6) providing evidence for this estimate.

### D.2. Comparison with LDA and NNMF

There are many methods for factorizing count data. The general goal of these models is to learn and low-dimensional representation of the high dimensional non-negative count data. Two popular methods are LDÄ (Blei, Ng and Jordan, 2003; Pritchard, Stephens and Donnelly, 2000) and NNMF (Lee and Seung, 1999).

#### Latent Dirichlet Allocation

Model hDP is related to LDA, but different in several critical aspects: (1) LDA is a Bayesian parametric model, whereas our model is a Bayesian nonparametric model, (2) LDA has one level of Dirichlet hierarchy, whereas our model has three, (3) standard LDA does not assume sample-specific subpopulation prior concentration, whereas our model integrates prior information about sample concentration. The last difference could be accommodated in the LDA model by assuming different Dirichlet parameter values, but we have not seen it done previously in practice. In our simulation experiments, we explore the performance of standard LDA and LDA incorporating varying prior parameters.

#### Non-negative matrix factorization

Non-negative matrix factorization is a natural method for factorizing read-count data because the read count matrix has all nonnegative entries. However, NNMF is very different than Model hDP. NNMF does not aim to estimate a distribution over subpopulations, does not allow for structured datasets, and does not allow one to specify the a priori subpopulation concentration for each sample. Nevertheless, because of the existence of fast inference algorithms, NNMF has been used for read-count datasets.

#### Results

To assess the importance of the hierarchical structure in Model hDP, we compared the performance to LDA and NNMF in terms of the KL divergence — lower values indicate the estimated distributions are closer to true distributions. Since both LDA and NNMF failed to find the true components in the simulation of mixture of bulk sample and single cell sample, we generated data sets using the same parametric model but only having bulk data and compared the performance with our model. Figure 11 shows a visualization of the comparisons between LDA, NNMF, and Model hDP. In Table 2, the values in the parentheses are the 95% confident interval of the KL divergence. Bold values are the best one in the same scenario. This experiment shows that Model hDP outperforms LDA and NNMF in all but one data scenario.

**Fig 10:**
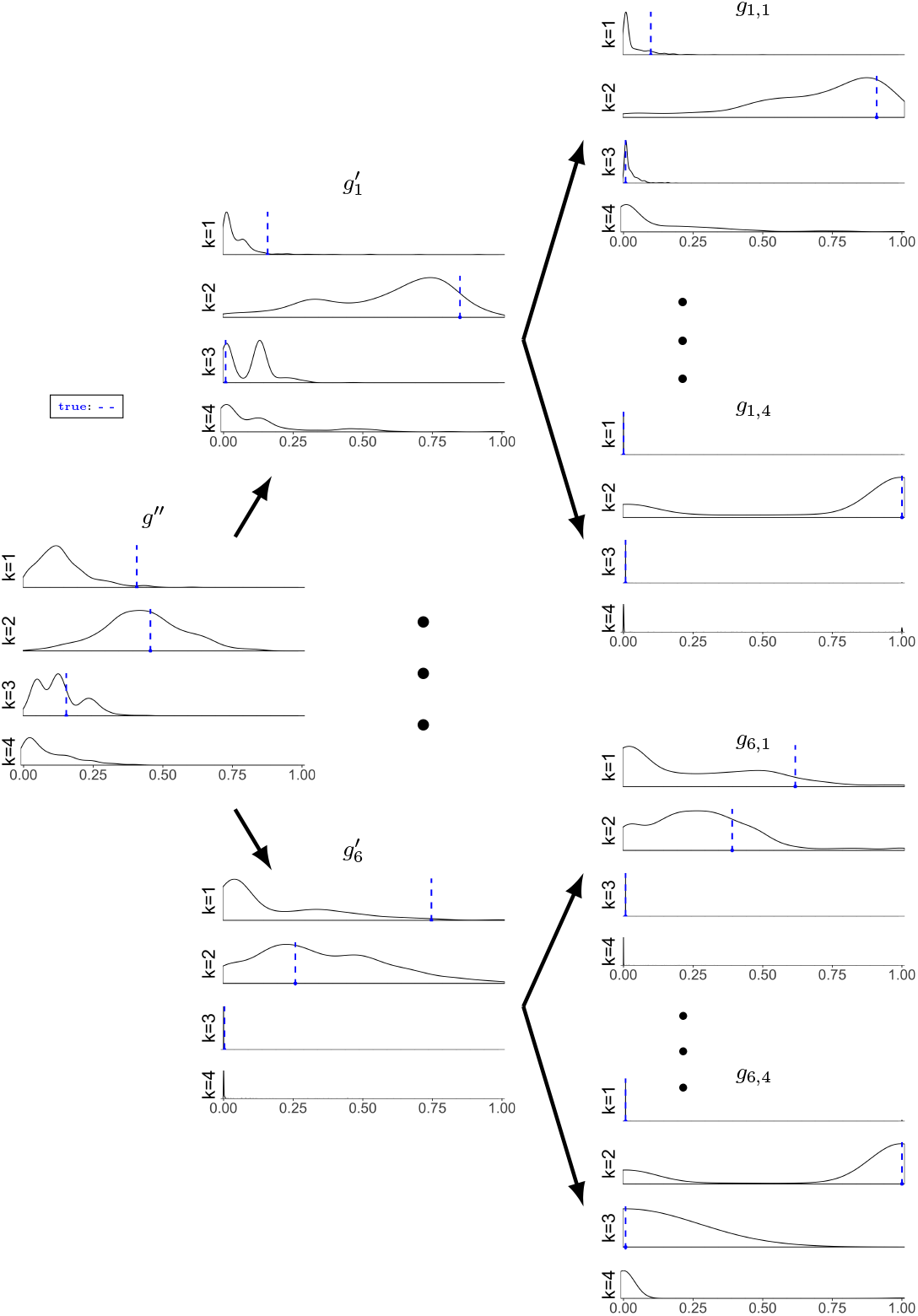
Marginal posterior density estimates for simulation data with *N* = 6 and *N_i_* =4

**Fig 11:**
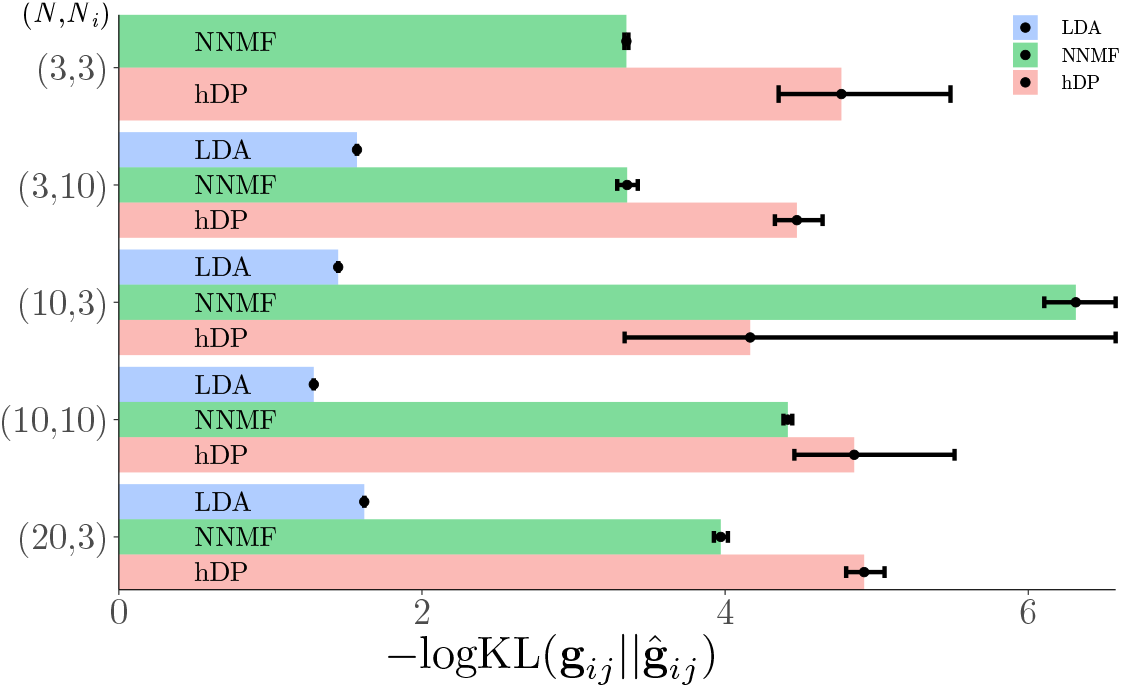
Comparisons among NNMF, LDA, and Model hDP. The KL divergence between the estimated sample-level posterior distribution over subpopulations and true distribution over subpopulations, 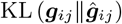, shows the model accuracy (higher — logKL is better).

### D.3. Comparison with Pyclone, PhyloWGS and TreeClone

#### Pyclone

When we analyzed the simulation data with Pyclone we found that it predicts all samples come from the same cluster with 0.558 prevalence (55.8% of the samples has the mutation) and 0.16 standard deviation. Pyclone does not have the capability to identify subpopulation frequencies above the sample level so we were not able to compare distributions at the individual and population level. Deshwar et al. (2015) noted that copy-number variation can be a powerful data modality for discriminating subpopulations with similar frequencies.

#### PhyloWGS

We attempted to analyze simulation data that was generated in the same way as describe in the *Data Generation* paragraph. We set the number of individuals, *N* = 5, the number of samples per individual, *N_i_* = 3, and number of reads per sample (across 5 genomic locations) to *R_ij_* = 100 — each genomic location has an average of 20 reads. We found that PhyloWGS was not able to produce enough posterior samples for inference. The method returned an error indicating that all samples are multiprimary—the posterior samples are polyclonal (too many components, thus not converged)—(https://github.com/morrislab/phylowgs/blob/master/pwgsresults/result_munger.py). So, we reduced the number of samples as indicated in the main text.

**Fig 12:**
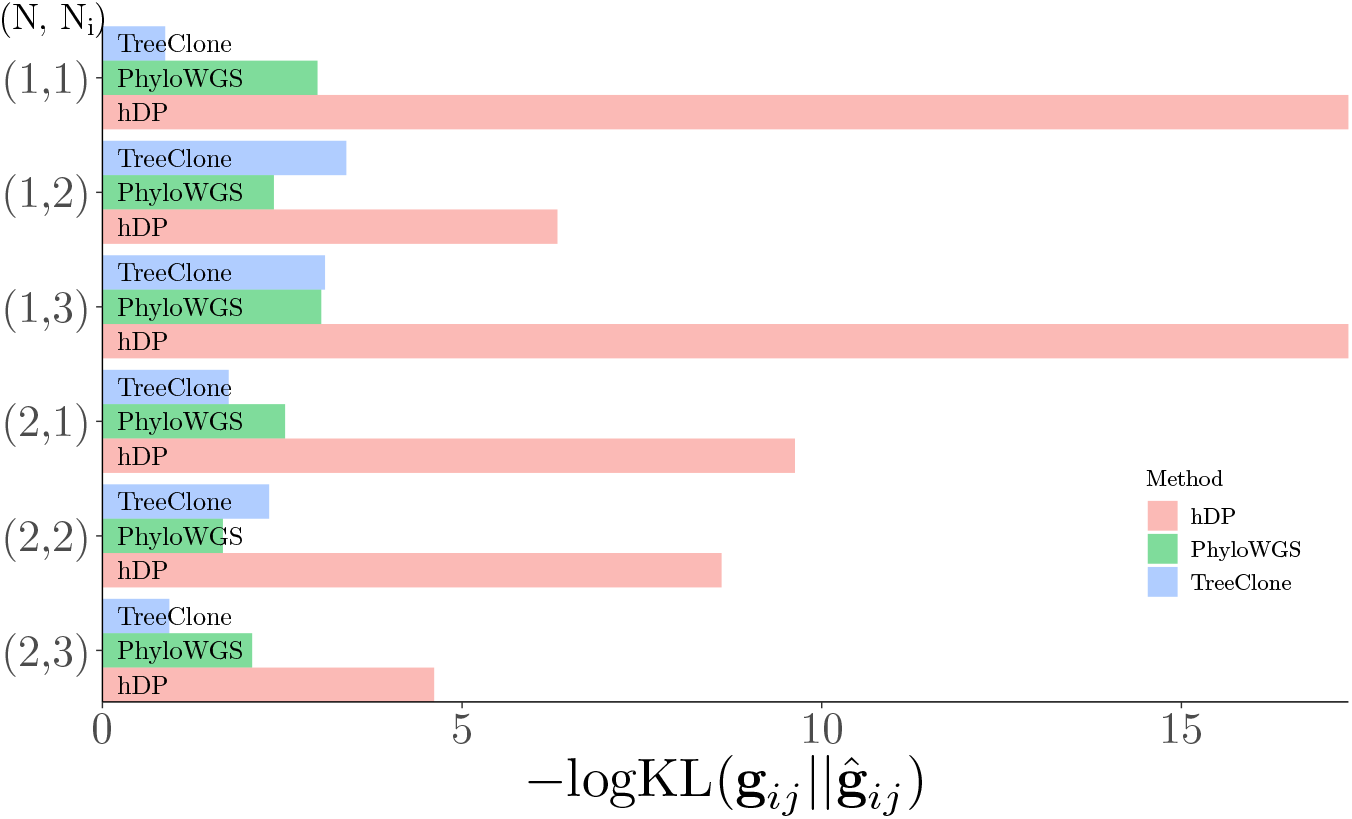
Comparison between Model hDP, PhyloWGS and TreeClone by KL divergence between estimated six sample-level posterior distribution and true distribution, 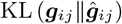, shows the model accuracy (higher — logKL is better).

## APPENDIX E: SENSITIVITY TO VARYING h, *K, L*, AND *ϵ*

To assess the sensitivity of the model, we performed simulation experiments varying 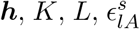, and 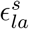. Data was generated as in Section 5. The number of individuals is *N* = 6, and the number of samples per individuals is *N_i_* = 4, where each individual has 1 bulk sample and 3 single-cell samples.

The metric used to assess the goodness-of-fit of the marginal posterior distribution of the subtypes is the standard KL divergence between the true posterior and the estimated posterior. The metric to assess the goodness-of-fit of the subpopulation genotype estimate is a modified Hamming distance:

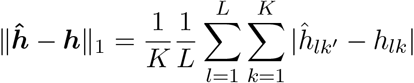

where *k*’ is subpopulation that is the closest match to the true genotype in the estimated genotype matrix, ***ĥ***.

### E.1. Sensitivity to varying *h*

We assessed the sensitivity of the model to variations in the genotype-subpopulation matrix ***h***. We used the same settings as the benchmark experiments: *K* = 3, *L* = 5, *α* = 1, *β_i_* = 1 for all 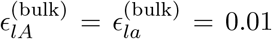 for bulk data and 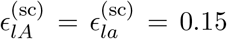 for single cell data. For the sample-level hyperparameters we set *γ_ij_* = 10 for for the bulk samples and *γ_ij_* = 0.1 for the single cell samples. We set the number of reads per sample (across 5 genomic locations), *R_ij_* = 100 — each genomic location has an average of 20 reads. The variables ***h*** was sampled randomly from *p*(*h_lk_* = (0,1, 2)) = (0.45, 0.1, 0.45). The marginal posterior distributions ***g***″|***x, g***′|***x, g***|***x*** and ***h*** were estimated using MCMC samples generated from Algorithm 1. We sampled 490 posterior samples after a burn-in/warm-up of 1,000 samples and thinning by a factor of 100. Table 3 shows the summary statistics for each of five random samples of ***h***.

**Table 3.** Simulation results of varying **h**. The values in parentheses of KL divergence columns are the standard deviation of the KL divergence.

| Experiments | $\ \hat{h} - h\ _1$ | $\text{KL}(g''\vert \vert \hat{g}'')$ | $\text{KL}(g'_i\vert \vert \hat{g}'_i)$ | $\text{KL}(g_{ij}\vert \vert \hat{g}_{ij})$ |
| --- | --- | --- | --- | --- |
| 1 | 0.133 | 1.122 | 2.046 (0.060) | 2.065 (1.10) |
| 2 | 0 | 1.435 | 1.682 (0.696) | 2.029 (1.215) |
| 3 | 0 | 1.720 | 2.296 (0.522) | 3.226 (1.942) |
| 4 | 0.133 | 1.008 | 1.596 (0.697) | 2.445 (1.816) |
| 5 | 0 | 0.917 | 1.683 (0.310) | 3.068 (1.980) |

From the table we can see the model successfully identifies the genotypes of all of the subpopulations exactly in 3 out of 5 experiments and makes only small errors in the two others. The marginal posterior distributions are close to the true distribution as well.

**Fig 13:**
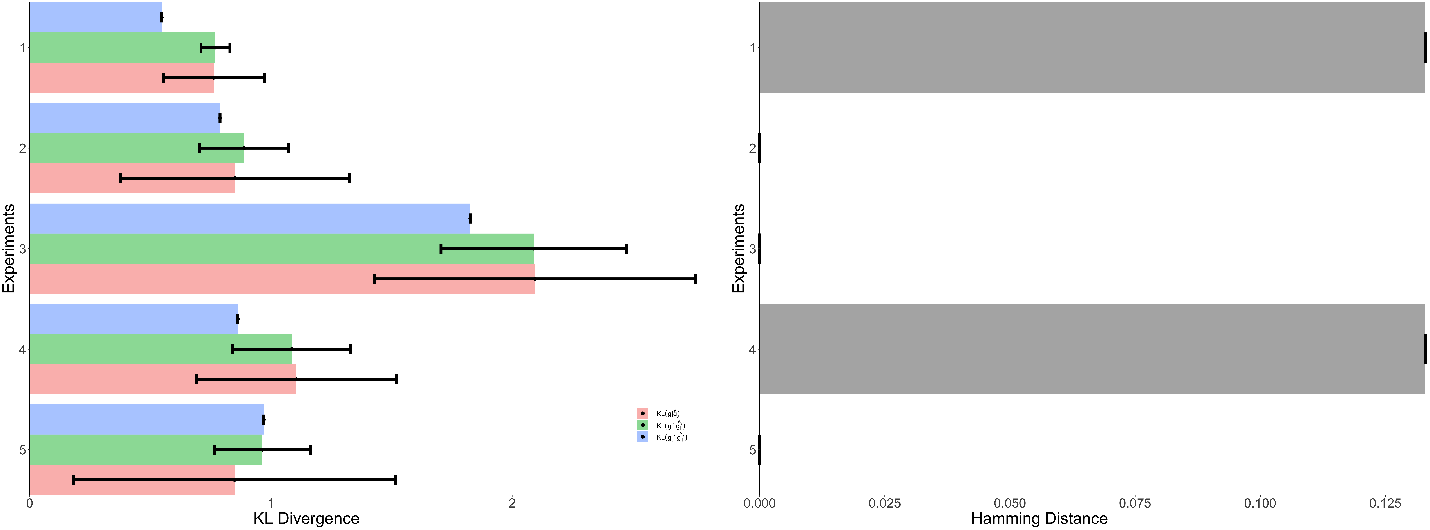
Sensitivity analysis varying ***h***. (left) Average KL divergence between true subpopulation distribution and estimated. (right) Average distance between true subpopulation genotype and estimated for five replicates.

### E.2. Sensitivity to varying *K*

Five groups of simulations with different *K* were conducted to examine the sensitivity of the model to varying the number of subpopulations, *K*. We set *K* ∈ {4, 5, 6, 8,10} for each group, and in each group we did three simulations with different realizations of ***h***. We set number of genomic locations, *L* = 10, to ensure there are is sufficent genotype space for different subpopulations when K is larger. we set number of reads per sample (across 10 genomic locations), *R_ij_* = 100 — each genomic location has an average of 10 reads.

We used the same settings as the benchmark experiments: *α* = 1, *β_i_* = 1 for all 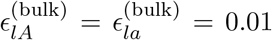 for bulk data and 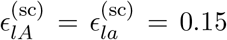 for single cell data. For the sample-level hyperparameters we set *γ_ij_* = 10 for for the bulk samples and *γ_ij_* =0.1 for the single cell samples. The matrix ***h*** was sampled with *p*(*h_lk_* = (0,1, 2)) = (0.45, 0.1, 0.45) randomly. The marginal posterior distributions ***g***″|***x, g***′|***x, g***|***x*** and ***h*** were estimated using MCMC samples generated from Algorithm 1. We sampled 490 posterior samples after a burn-in/warm-up of 1,000 samples and thinning by a factor of 100. Whether the model finds all true components and the mean KL divergence at each level for five experiments are shown below in Table 4 and Figure 14.

**Fig 14:**
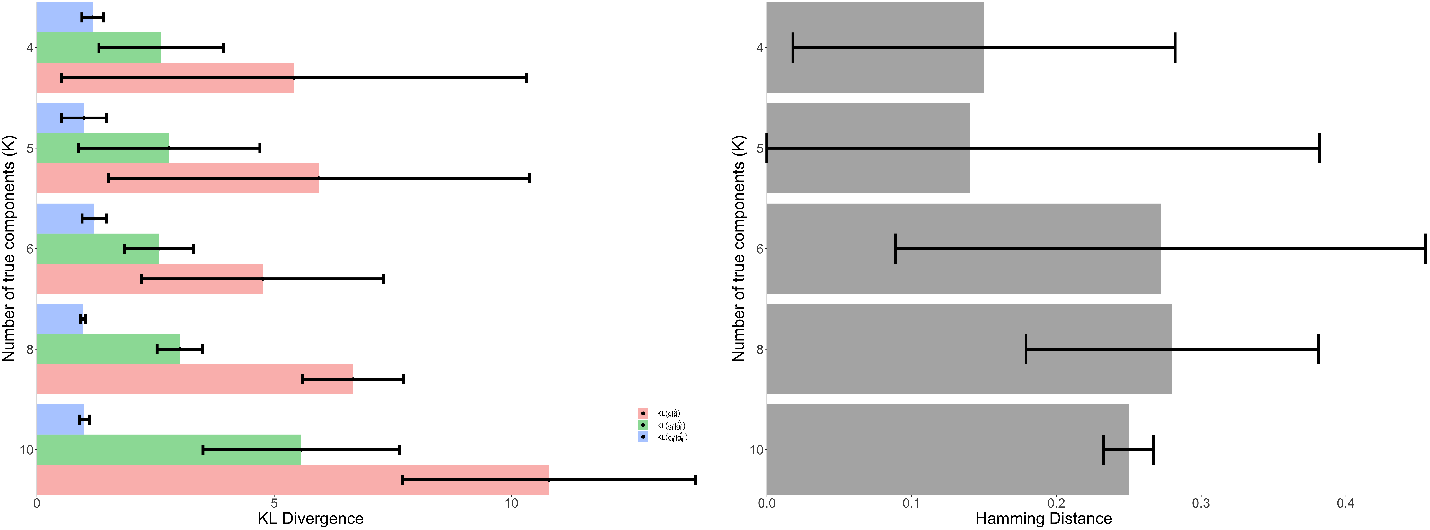
Sensitivity analysis varying *K*. (left) Average KL divergence between true subpopulation distribution and estimated. (right) Average distance between true subpopulation genotype and estimated.

**Table 4.** Simulation results of varying K. The values in parentheses of KL divergence columns are the standard deviation of the KL divergence.

| $K$ | $\ \hat{\mathbf{h}} - \mathbf{h}\ _1$ | $\text{KL}(\mathbf{g}'' \ \hat{\mathbf{g}}'')$ | $\text{KL}(\mathbf{g}'_i \ \hat{\mathbf{g}}'_i)$ | $\text{KL}(\mathbf{g}_{ij} \ \hat{\mathbf{g}}_{ij})$ |
| --- | --- | --- | --- | --- |
| 4 | 0.15(0.132) | 1.171 (0.224) | 2.619 (1.312) | 5.419 (4.90) |
| 5 | 0.14(0.242) | 0.994 (0.473) | 2.785 (1.914) | 5.941 (4.438) |
| 6 | 0.272(0.183) | 1.206 (0).255 | 2.576 (0.722) | 4.760 (2.551) |
| 8 | 0.280(0.101) | 0.969 (0.049) | 3.010 (0.479) | 6.664 (1.062) |
| 10 | 0.25(0.017) | 1 (0.110) | 5.574 (2.074) | 10.793 (3.086) |

From the table we can see, even the number of true component increased 10, the model can find the true components of ***h***. The main reason that some simulations can’t find the remaining components is that the parametric model does not generate enough remaining component data at sample level.

### E.2. Sensitivity to varying *L*

Five groups of simulations with different values of *L* were conducted to examine the sensitivity of L. We set *L* ∈ {5,10, 20, 50,100} for each group, and in each group we did three simulations with randomly sampled ***h***. The true number of subpopulations was set to *K* = 3

We kept other settings same as we did in the previous section: *α* = 1, *β_i_* = 1 for all 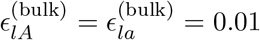 for bulk data and 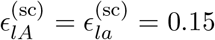 for single cell data. The prior concentration parameter for each sample was set to *γ_ij_* = 10 for bulk samples and *γ_ij_* = 0.1 for single-cell samples. The subpopulation-genotype matrix ***h*** was generated with a prior where *p*(*h_lk_* = (0,1, 2)) = (0.45, 0.1, 0.45) randomly. The marginal posterior distributions ***g***″|***x, g***‱|***x, g***|***x*** and ***h*** were estimated using MCMC samples generated from Algorithm 1. To reduce the running time, we sampled 900 posterior samples after a burn-in/warm-up of 1,000 samples and thinning by a factor of 10.

Table 5 and Figure 15 show the accuracy of the estimated distribution over subpopulations and subpopulation genotypes according to the metrics previously defined. The results show that the goodness-of-fit and accuracy of the products of inference are stable for a wide range of the number of targeted genomic loci.

**Fig 15:**
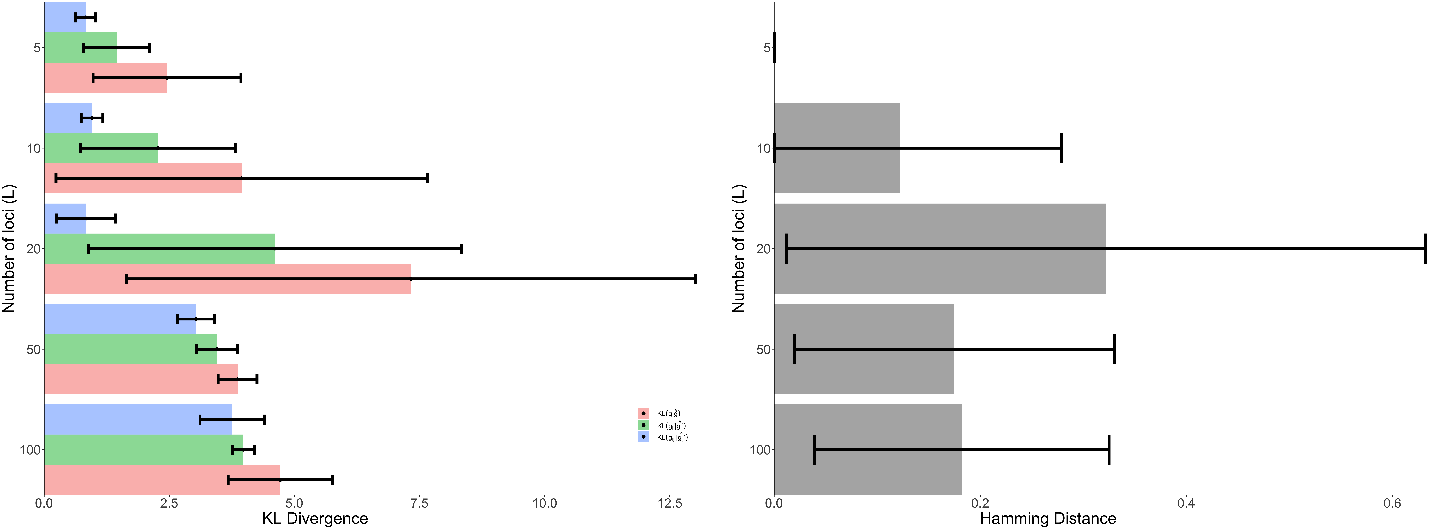
Sensitivity analysis varying *L*. (left) Average KL divergence between true subpopulation distribution and estimated. (right) Average distance between true subpopulation genotype and estimated.

**Table 5.** Simulation results of varying L. The values in parentheses of KL divergence columns are the standard deviation of the KL divergence.

| L | $\ \hat{\mathbf{h}} - \mathbf{h}\ _1$ | $\text{KL}(\mathbf{g}''\vert \mathbf{g}'')$ | $\text{KL}(\mathbf{g}'\vert \mathbf{g}')$ | $\text{KL}(\mathbf{g}_{ij}\vert \mathbf{g}_{ij})$ |
| --- | --- | --- | --- | --- |
| 5 | 0 | 0.892 (0.201) | 1.448 (0.660) | 2.455 (1.477) |
| 10 | 0.122 (0.157) | 0.957 (0.217) | 2.27 (1.547) | 3.944 (3.708) |
| 20 | 0.322 (0.310) | 0.837 (0.585) | 4.611 (3.722) | 7.327 (5.685) |
| 50 | 0.175 (0.154) | 3.032 (0.369) | 3.454 (0.414) | 3.864 (0.388) |
| 100 | 0.182 (0.143) | 3.754 (0.640) | 3.978 (0.220) | 4.720 (1.046) |

### E.4. Sensitivity to varying *ϵ*

In the real data analysis, specifying the sequencing error parameter for single-cell data can be problematic, because in practice, the error is unknown (although for bulk data it can be argued the error is small). Thus we conduct five groups of simulations to examine the sensitivity of 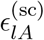 and 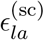. We set 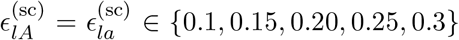. For each group, and in each group we did three simulations with different values of ***h*** and we set the number of true subpopulations to *K* = 3

We kept other settings same as we did in the previous section: *α* = 1, *β_i_* = 1 for all *i*. The subpopulation-genotype matrix ***h*** was generated with a prior where *p*(*h_lk_* = (0,1, 2)) = (0.45, 0.1, 0.45) randomly. The marginal posterior distributions ***g***″|***x, g***′|***x, g***|***x*** and ***h*** were estimated using MCMC samples generated from Algorithm 1. We sampled 490 posterior samples after a burn-in/warm-up of 1,000 samples and thinning by a factor of 100.

Table 6 and Figure 16 show the accuracy of the estimated distribution over subpopulations and subpopulation genotypes according to the metrics previously defined. There is a slight degradation of performance of the subpopulation distribution estimates as the number error rate of the single-cell data is increased. We conjecture that the robustness is a product of the constraint that the -subpopulation genotype matrix must be discrete and multiple simultaneous sequencing errors would be required to overwhelm the capability of the model to infer the discrete genotype.

**Fig 16:**
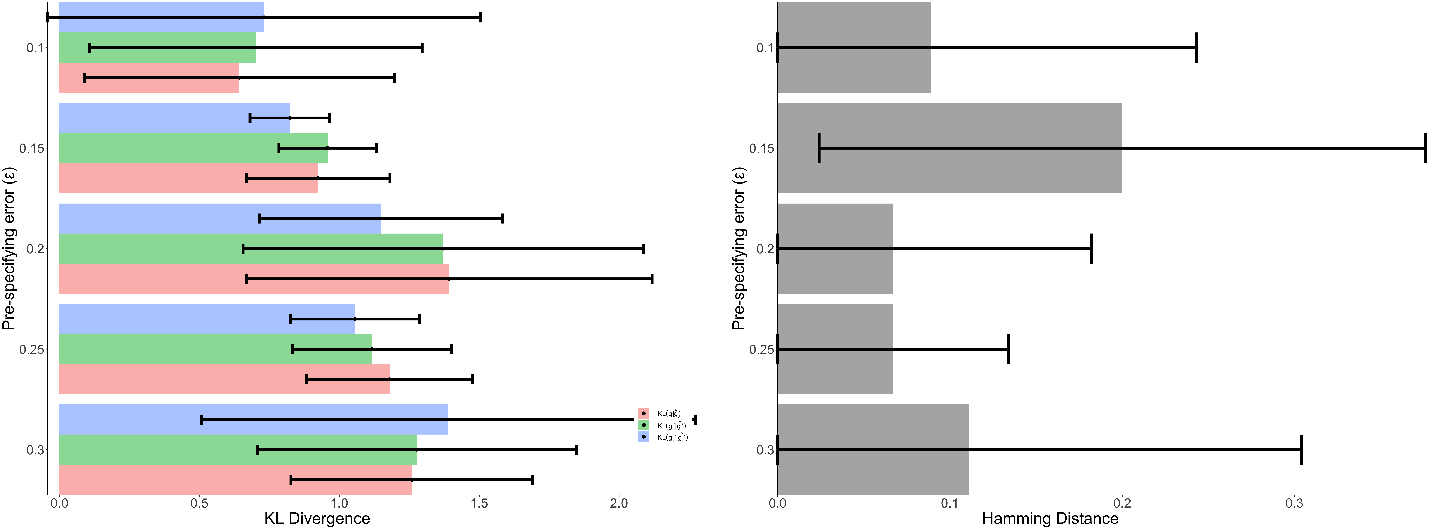
Sensitivity analysis varying 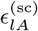 and 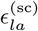. (left) Average KL divergence between true subpopulation distribution and estimated. (right) Average distance between true subpopulation genotype and estimated.

**Table 6.** Simulation results of varying 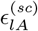 and, 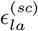. The values in parentheses of KL divergence columns are the standard deviation of the KL divergence.

| $\epsilon_{lA}^{(sc)} = \epsilon_{la}^{(sc)}$ | $\ \hat{\mathbf{h}} - \mathbf{h}\ _1$ | $\text{KL}(\mathbf{g}''\ \hat{\mathbf{g}}'')$ | $\text{KL}(\mathbf{g}'\ \hat{\mathbf{g}}'_i)$ | $\text{KL}(\mathbf{g}_{ij}\ \hat{\mathbf{g}}_{ij})$ |
| --- | --- | --- | --- | --- |
| 0.10 | 0.089 (0.154) | 0.731 (0.773) | 0.701 (0.595) | 0.642 (0.555) |
| 0.15 | 0.200 (0.176) | 0.823 (0.142) | 0.958 (0.174) | 0.924 (0.257) |
| 0.20 | 0.067 (0.115) | 1.149 (0.435) | 1.371 (0.715) | 1.393 (0.724) |
| 0.25 | 0.067 (0.067) | 1.056 (0.231) | 1.116 (0.284) | 1.180 (0.297) |
| 0.3 | 0.111 (0.192) | 1.39 (0.883) | 1.277 (0.570) | 1.259 (0.432) |

## APPENDIX F: ALL DATASET ANALYSIS

### F.1. Genomic Loci Selected for Inference

Inference on the ALL dataset using Algorithm 1 used a curated subset of loci from the original study paper (Gawad, Koh and Quake, 2014). Table 7 shows a listing of all 111 nonsynonymous mutations identified in that study referenced to hg38 coordinates. A star next to the locus indicates it was selected in the curated subset. Inference on the ALL dataset using Algorithm 2 used the full set of 111 loci of which 109 are identifiable in the full set of single-cell data.

**Table 7.** Full set of 111 Loci in hg38 coordinates

| Locus (hg38) | Gene | Locus (hg38) | Gene | Locus (hg38) | Gene |
| --- | --- | --- | --- | --- | --- |
| chr1:36282610 | THRAP3 | chr8:25319665 | DOCK5 | chr17:10523184 | MYH2 |
| chr1:41484129 | EDN2 | chr8:42305190 | IKBKB | chr17:39439170 | MED1 |
| chr1:47144592 | CYP4A22 | chr8:73081054 | SBSPON | chr17:45241470 | FMNL1 |
| chr1:65429930 | LEPROT | chr8:104013561 | RIMS2 | chr17:58087167 | DYNLL2 |
| chr1:74572702 | C1orf173 | chr8:109119136 | TRHR | chr17:58327917 | BZRAP1 |
| chr1:108699693 | PRPF38B | chr8:125011548 | SQLE | chr18:69599492 | DOK6 |
| chr1:220136888 | IARS2 | chr8:143468539 | ZC3H3 | chr19:10291260 | ICAM5 |
| chr2:36743187 | VIT | chr8:143838738 | NRBP2 | chr19:12764849 | HOOK2 |
| chr2:51028023 | NRXN1 | chr8:143920874* | PLEC | chr19:19349426 | MAU2 |
| chr2:102340723 | IL1RL1 | chr8:143924816* | PLEC | chr19:37612716 | ZNF540 |
| chr2:108749028 | RANBP2 | chr9:89605491 | GADD45G | chr19:37669839 | ZNF781 |
| chr2:169637471* | PPIG | chr9:113596819 | RGS3 | chr19:40389761 | HIPK4 |
| chr2:178531094 | TTN | chr9:114406547 | DFNB31 | chr19:44819680 | BCAM |
| chr2:178751268 | TTN | chr9:130053958 | GPR107 | chr19:51414455 | SIGLEC10 |
| chr2:191846921 | SDPR | chr9:137028806 | ABCA2 | chr19:52384775* | ZNF880 |
| chr2:195853387 | DNAH7 | chr10:32293720 | EPC1 | chr20:45222995 | SEMG2 |
| chr2:219575504 | INHA | chr10:69097200 | SRGN | chr20:53253823 | TSHZ2 |
| chr2:231108908 | HTR2B | chr10:100295080 | PKD2L1 | chr21:46112405 | COL6A2 |
| chr3:51225696 | DOCK3 | chr10:100924346* | FAM178A | chr22:37643750 | SH3BP1 |
| chr3:147413487 | ZIC1 | chr10:100924409* | FAM178A | chrX:20042572 | MAP7D2 |
| chr3:149207404 | CP | chr10:116555216 | PNLIP | chrX:34944563 | FAM47B |
| chr3:160279300 | IFT80 | chr10:117137896 | VAX1 | chrX:48802875 | HDAC6 |
| chr4:4274574 | LYAR | chr10:127374104 | DOCK1 | chrX:51895445 | MAGED1 |
| chr4:10077821 | WDR1 | chr11:6601517 | RRP8 | chrX:71141950 | MED12 |
| chr4:13542233 | NKX3-2 | chr11:71549007 | KRTAP5-9 | chrX:80677205 | BRWD3 |
| chr4:15007408 | CPEB2 | chr11:121550594 | SORL1 | chrX:102603414 | ARMCX5 |
| chr4:39874390 | PDS5A | chr12:25227337* | KRAS | chrX:118628217 | DOCK11 |
| chr4:147542558 | EDNRA | chr12:25245328* | KRAS | chrX:126165005 | DCAF12L2 |
| chr4:186618077* | FAT1 | chr12:80619462 | PTPRQ | chrX:141879215 | MAGEC3 |
| chr5:58458999 | PLK2 | chr12:104082836 | HCFC2 |  |  |
| chr5:81343827 | ACOT12 | chr12:107618313 | BTBD11 |  |  |
| chr5:139317101 | MATR3 | chr13:102746379 | CCDC168 |  |  |
| chr5:141399025 | PCDHGB5 | chr14:41887271 | LRFN5 |  |  |
| chr6:33799111 | MLN | chr14:69952279 | SMOC1 |  |  |
| chr6:56466116 | DST | chr15:33660305 | RYR3 |  |  |
| chr6:56476181 | DST | chr15:71960041 | MYO9A |  |  |
| chr6:131637429 | ENPP3 | chr16:66963323 | CES3 |  |  |
| chr7:12383545 | VWDE | chr16:67297619 | KCTD19 |  |  |
| chr7:18644770* | HDAC9 | chr16:70371413 | DDX19A |  |  |
| chr7:98978259 | TRRAP | chr17:1754190 | SERPINF2 |  |  |
| chr7:138906974 | KIAA1549 | chr17:7501644 | POLR2A |  |  |

### F.2. Samples selected for Inference

Inference on the ALL dataset using Algorithm 1 and Algorithm 2 used a subset of samples from the original study paper (Gawad, Koh and Quake, 2014). Table 8 shows a listing of the samples.

**Table 8.** *Samples selected from* ALL *data for data analysis.*

| Patient | Sample | Type | Read Count Across Ten Loci |
| --- | --- | --- | --- |
| 1 | Bulk | bulk | 632 |
|  | Cell_1_S10 | single-cell | 15 |
|  | Cell_1_S100 | single-cell | 11 |
|  | Cell_1_S101 | single-cell | 12 |
| 2 | Bulk | bulk | 792 |
|  | Cell_2_S10 | single-cell | 10 |
|  | Cell_2_S100 | single-cell | 18 |
|  | Cell_2_S101 | single-cell | 2 |
| 3 | Bulk | bulk | 806 |
|  | Cell_3_S100 | single-cell | 36 |
|  | Cell_3_S101 | single-cell | 45 |
|  | Cell_3_S118 | single-cell | 37 |
| 4 | Bulk | bulk | 850 |
|  | Cell_4_S101 | single-cell | 36 |
|  | Cell_4_S107 | single-cell | 32 |
|  | Cell_4_S110 | single-cell | 31 |
| 5 | Bulk | bulk | 837 |
|  | Cell_5_S10 | single-cell | 43 |
|  | Cell_5_S100 | single-cell | 51 |
|  | Cell_5_S101 | single-cell | 60 |
| 6 | Bulk | bulk | 873 |
|  | Cell_6_S10 | single-cell | 7 |
|  | Cell_6_S100 | single-cell | 9 |
|  | Cell_6_S101 | single-cell | 13 |

### F.3. Convergence

We used Geweke’s diagnostics to check the converge. Geweke’s diagnostics is the test that comparing mean of first 10% and last 50% of the MCMC chains (Geweke, 1991).

We used Geweke’s diagnostics function in pymc3 package to apply this test (John Salvatier Thomas V. Wiecki, 2016). This function compares mean of the first 10% samples of the chain and slices of the last 50 % samples of the chain and returns Z scores. Scores for a converged chain would oscillate between −1 and 1. Chains associated components with high probabilities passed the test. Figure 17 shows the scores of three chains of the Geweke’s diagnostics over different levels of some dimensions. We can see the scores are oscillating between −1 and 1.

**Fig 17:**
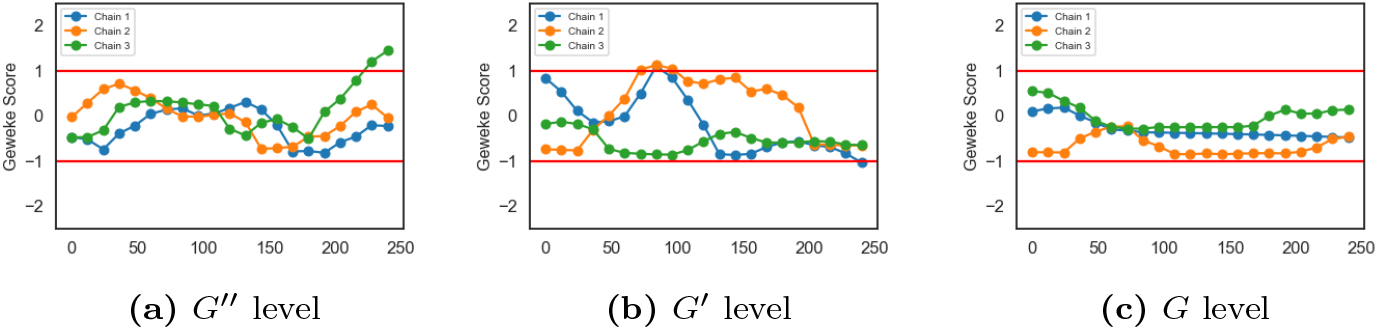
Geweke’s test at different levels

Trace plots in Figure 18 show that the sampler has converged as well.

**Fig 18:**
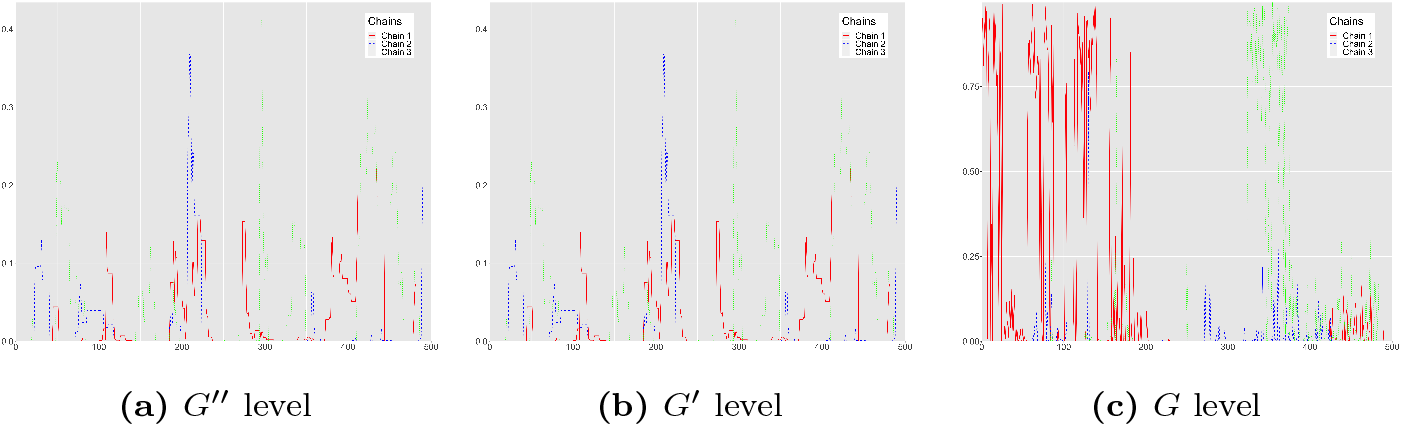
Trace plots at different levels of the model hierarchy.

### F.4. Read count tables and posterior distributions for Model hDP

The figures below are the combination of read count tables and posterior distributions for Patient 1-6. The left side are the read count tables for Patient 1-6 across ten loci (one bulk sample and three single-cell samples selected for each patient). The major/minor allele ratios are shown in parenthesis after each read count. Zero read counts are shown as dashes indicating missing data at those loci. The right side are the posterior distributions for Patient 1-6 at all levels. Red bars show the population level distribution over subpopulations 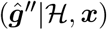, blue bars show the individual level distribution 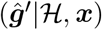, and green bars show the sample level distributions 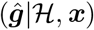, where 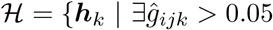, for *i* = 1, 2, 3, 4, 5, 6 and *j* = 1, 2, 3,4}.

### F.5. Posterior Distribution of h from Model hDP

Table 15 shows one posterior sample of **h. h** is discrete and each row represents a potential component which will associate with probabilities of 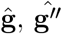, and 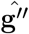.

**Table 9:** Read Count Table for Patient 1

|  | Patient 1 |  |  |  |
| --- | --- | --- | --- | --- |
|  | BULK | S10 | S100 | S101 |
| PPIG (chr2:169637471) | 52 (44/8) | 15 (13/2) | 11 (1/10) | 12 (12/0) |
| FAT1 (chr4:186618077) | 37 (36/1) | — | — | — |
| HDAC9 (chr7:18644770) | 56 (56/0) | — | — | — |
| PLEC (chr8:143920874) | 47 (47/0) | — | — | — |
| PLEC (chr8:143924816) | 90 (90/0) | — | — | — |
| FAM178A (chr10:100924346) | 72 (71/1) | — | — | — |
| FAM178A (chr10:100924409) | 30 (29/1) | — | — | — |
| KRAS (chr12:25227337) | 101 (101/0) | — | — | — |
| KRAS (chr12:25245328) | 124 (124/0) | — | — | — |
| ZNF880 (chr19:52384775) | 23 (23/0) | — | — | — |

**Table 10:**
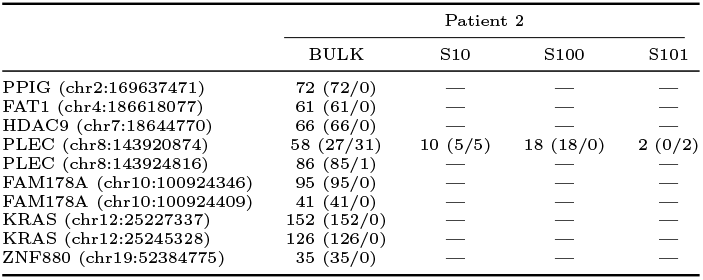
Read Count Table for Patient 2

**Table 11:**
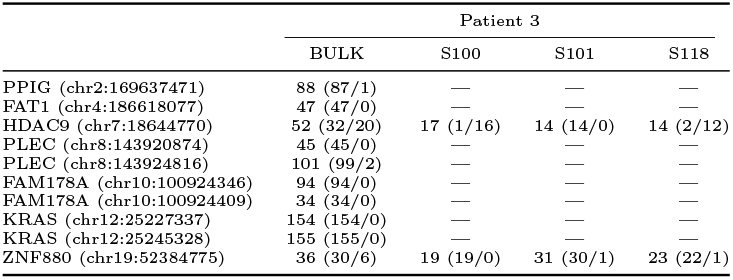
Read Count Table for Patient 3

**Table 12:**
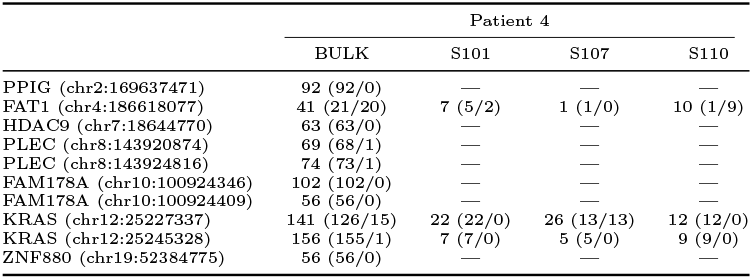
Read Count Table for Patient 4

**Table 13:**
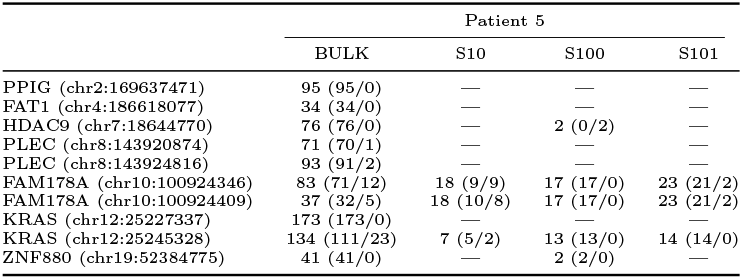
Read Count Table for Patient 5

**Table 14:**
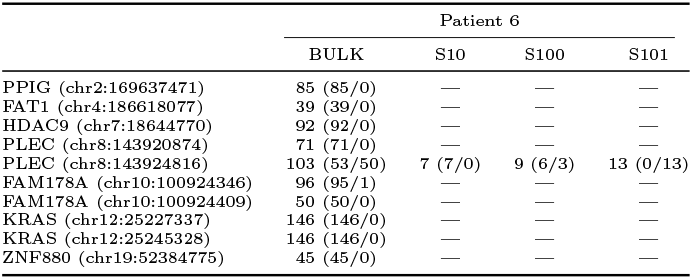
Read Count Table for Patient 6

**Table 15.**
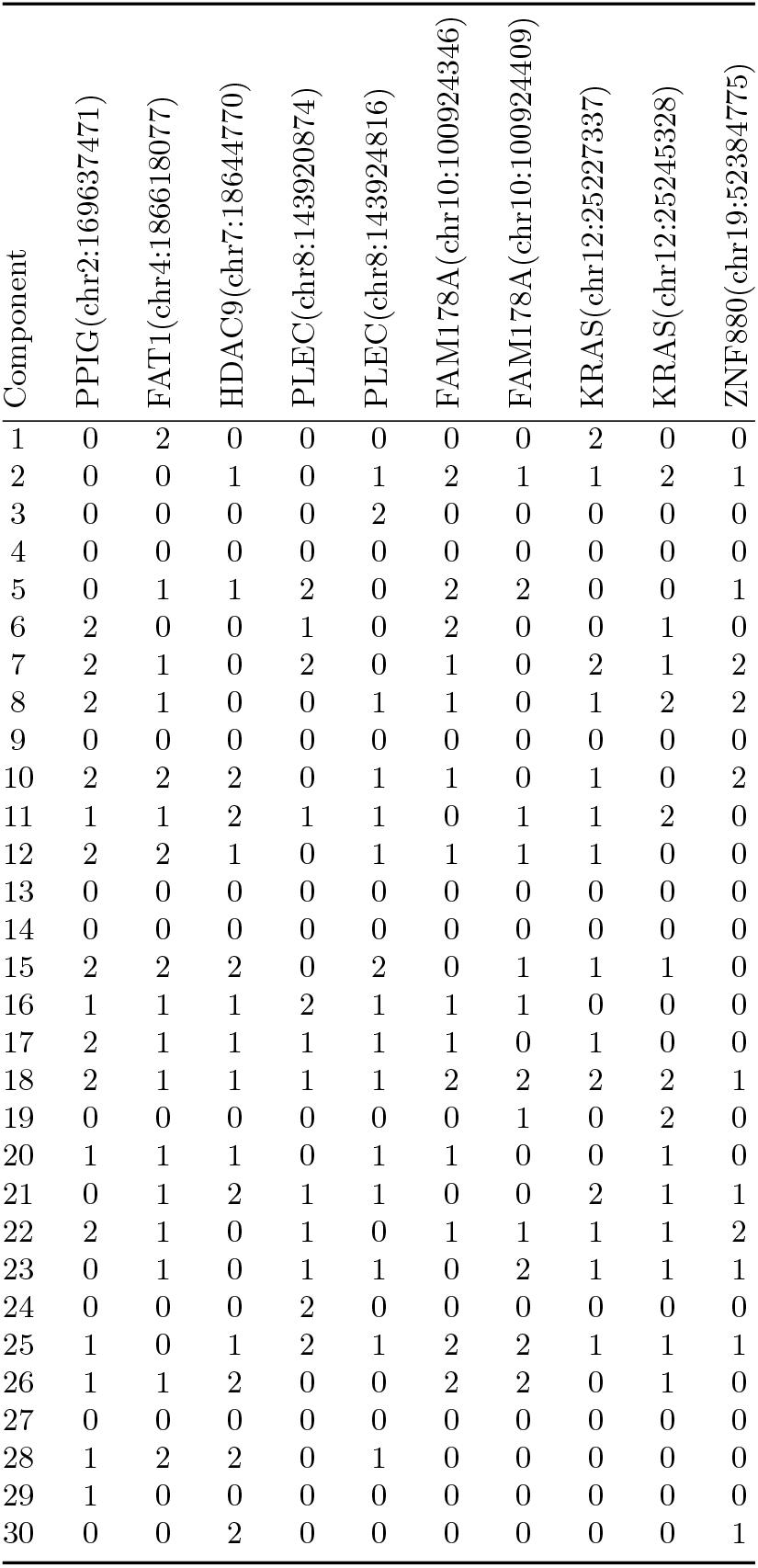
One posterior sample of **h** matrix, loci are shown in hg38 coordinates

### F.6. h matrix Inference from Gamma-Poisson Model

Figure 25 shows a posterior sample of **h** for the subpopulations identified for Patient 1.

**Fig 19:**
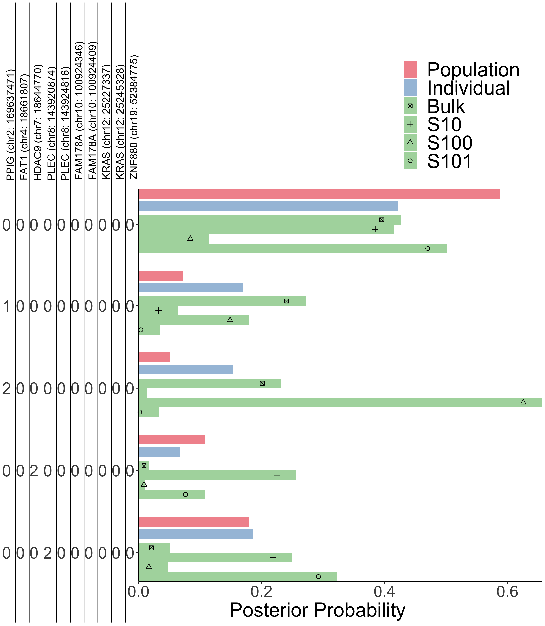
Posterior Distribution of Patient 1

**Fig 20:**
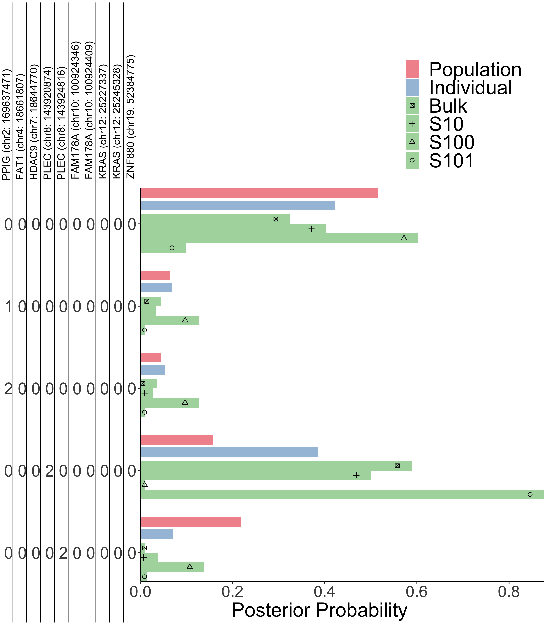
Posterior Distribution of Patient 2

**Fig 21:**
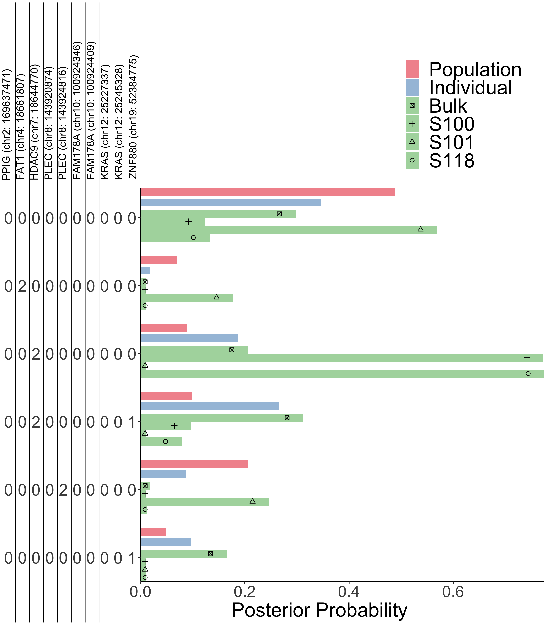
Posterior Distribution of Patient 3

**Fig 22:**
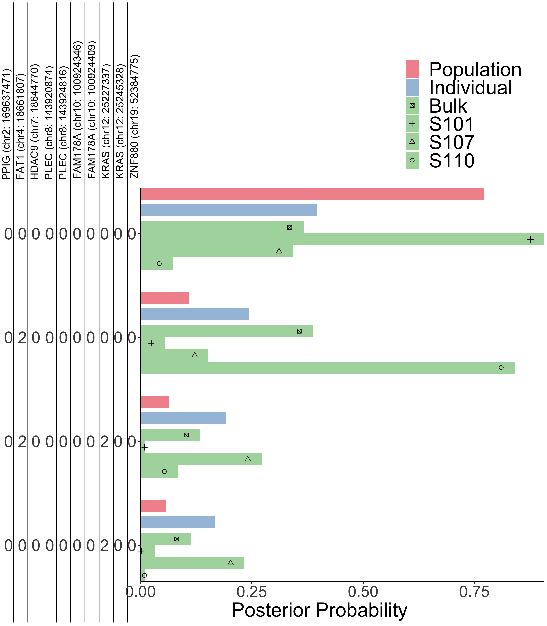
Posterior Distribution of Patient 4

**Fig 23:**
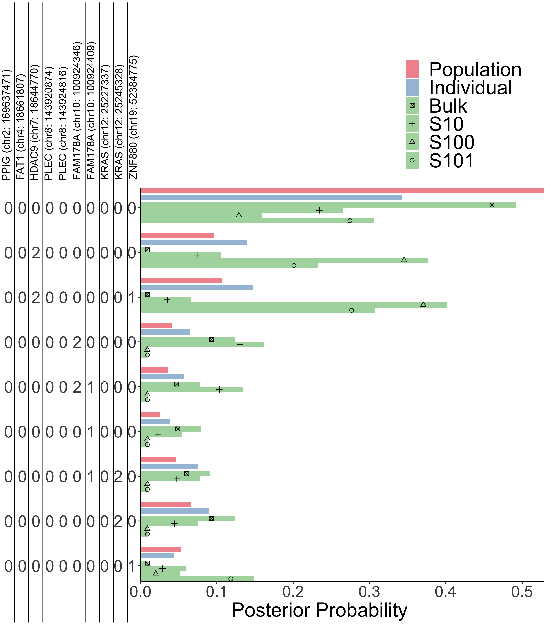
Posterior Distribution of Patient 5

**Fig 24:**
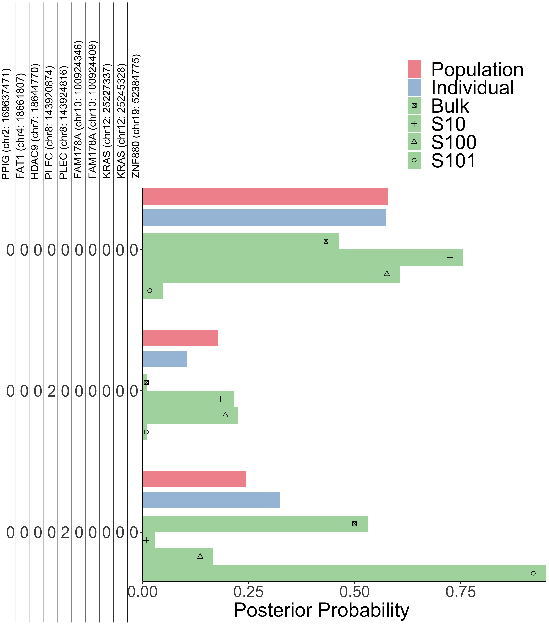
Posterior Distribution of Patient 6

**Fig 25:**
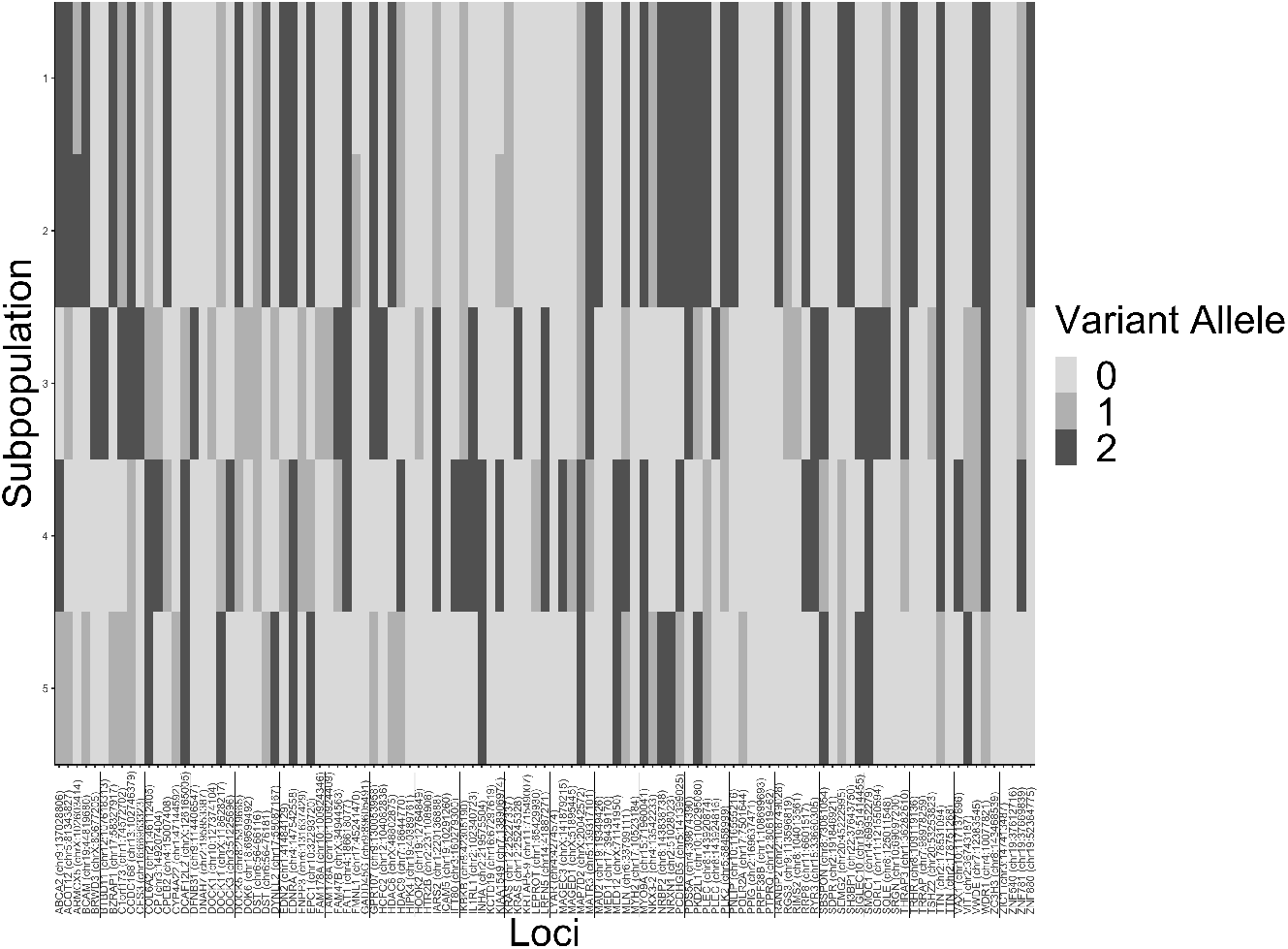
Posterior sample from **h** for Patient 1.

### F.7. Co-occurence Networks for Patients 1—6

The figures show the adjacency matrices in network form for Patient 1-6 where an edge between *l* and *l*’ is drawn if *a_ll′_* > 0.50. Loci without edges to other loci are omitted.

710 N. Pleasant St, Data Science Institute, Columbia University

Amherst, MA 01003

**Fig 26:**
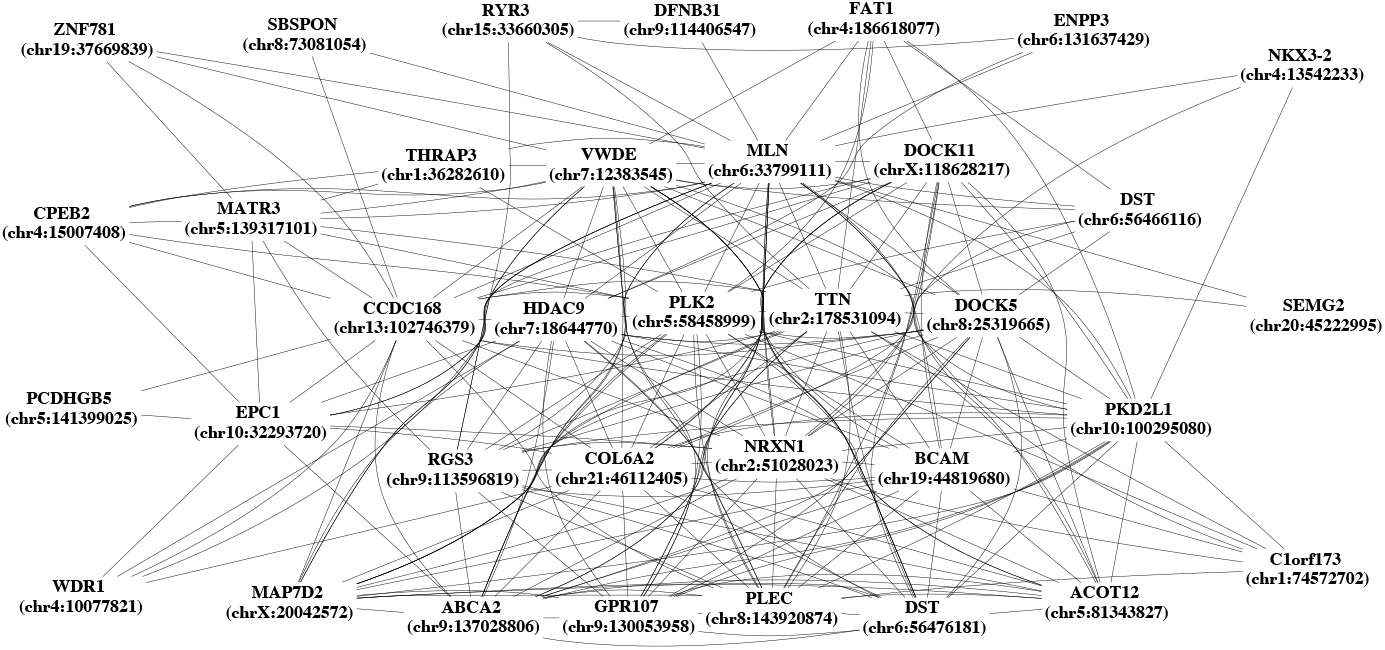
Inferred mutation co-occurence network across Patient 1 from Model hGP.

**Fig 27:**
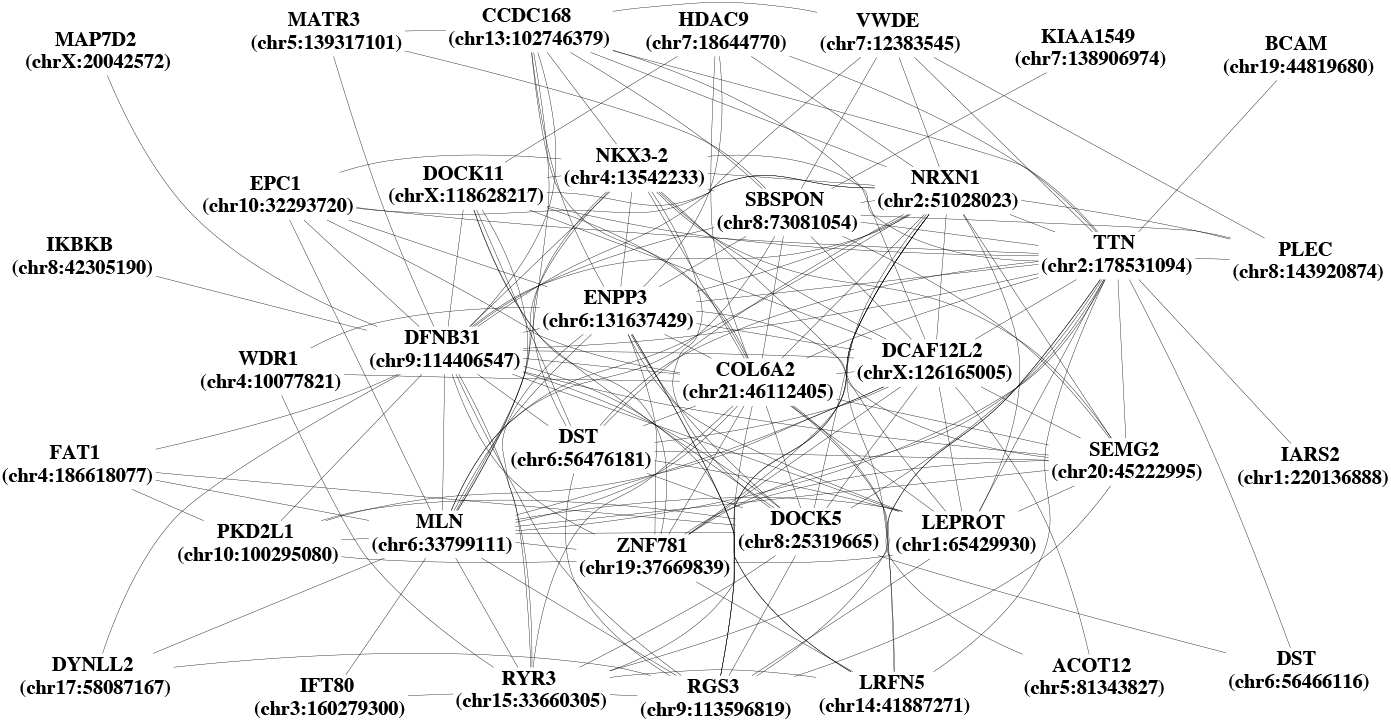
Inferred mutation co-occurence network across Patient 2 from Model hGP.

**Fig 28:**
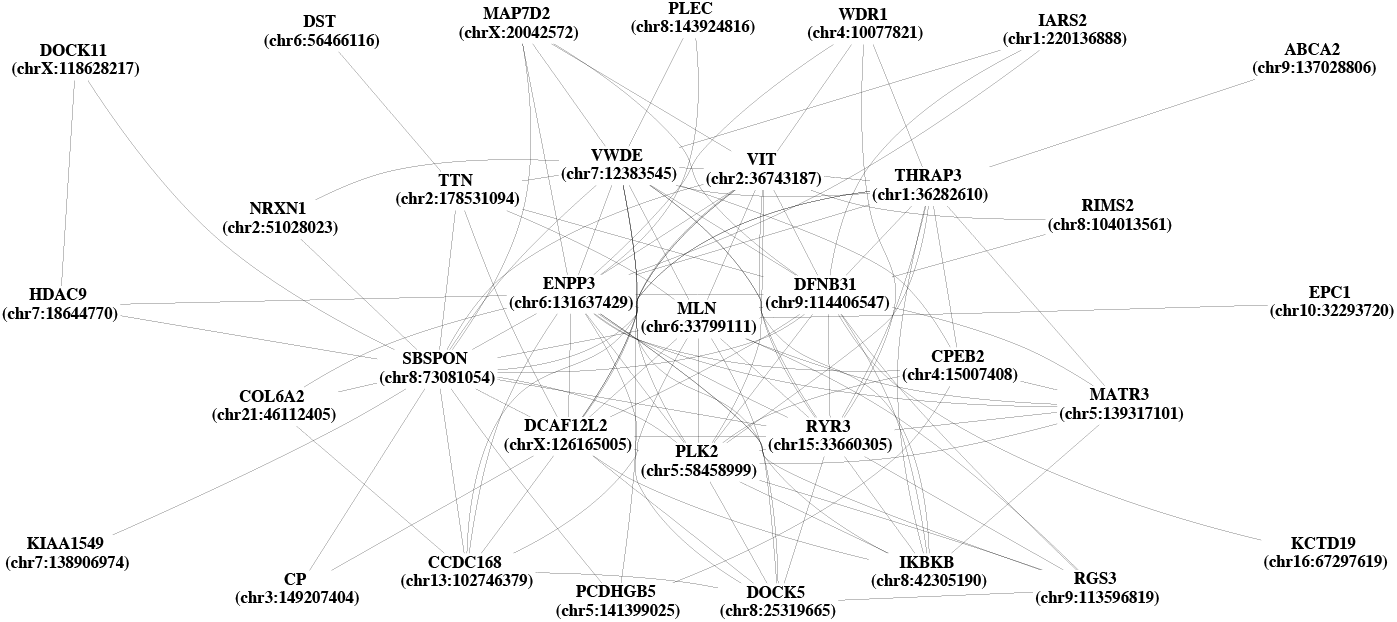
Inferred mutation co-occurence network across Patient 3 from Model hGP.

**Fig 29:**
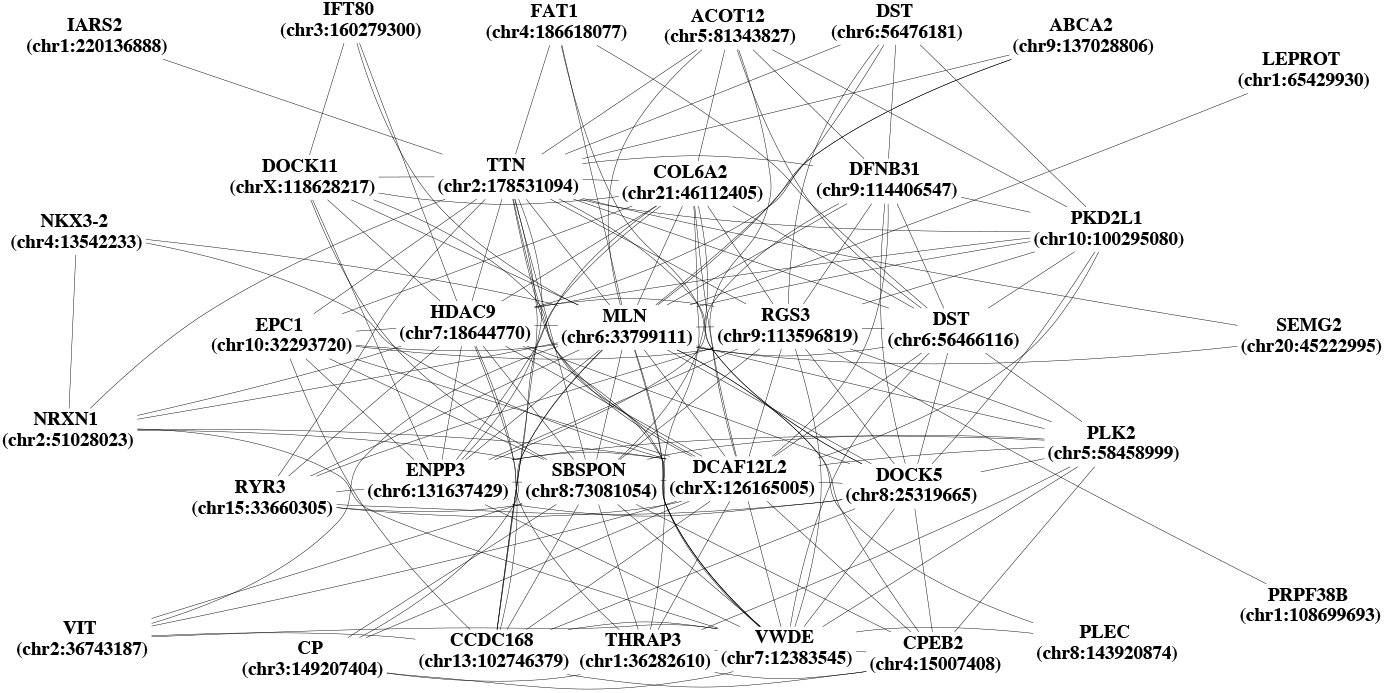
Inferred mutation co-occurence network across Patient 4 from Model hGP.

**Fig 30:**
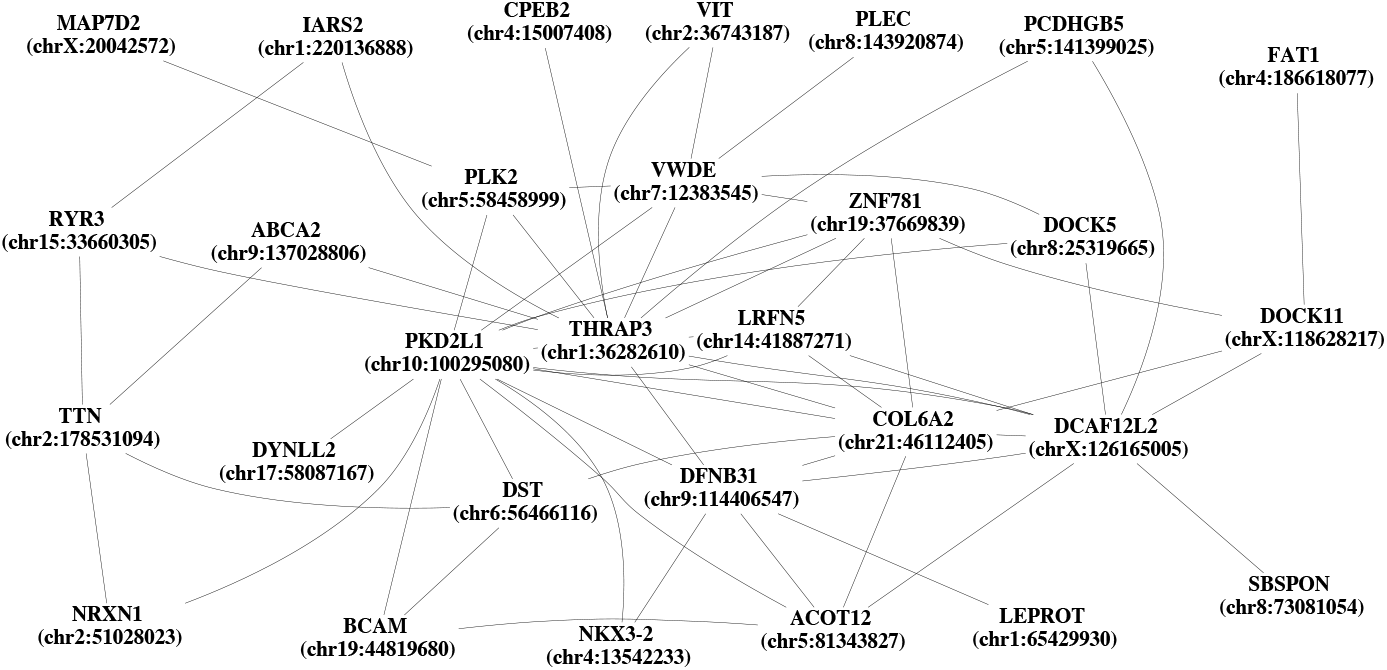
Inferred mutation co-occurence network across Patient 5 from Model hGP.

**Fig 31:**
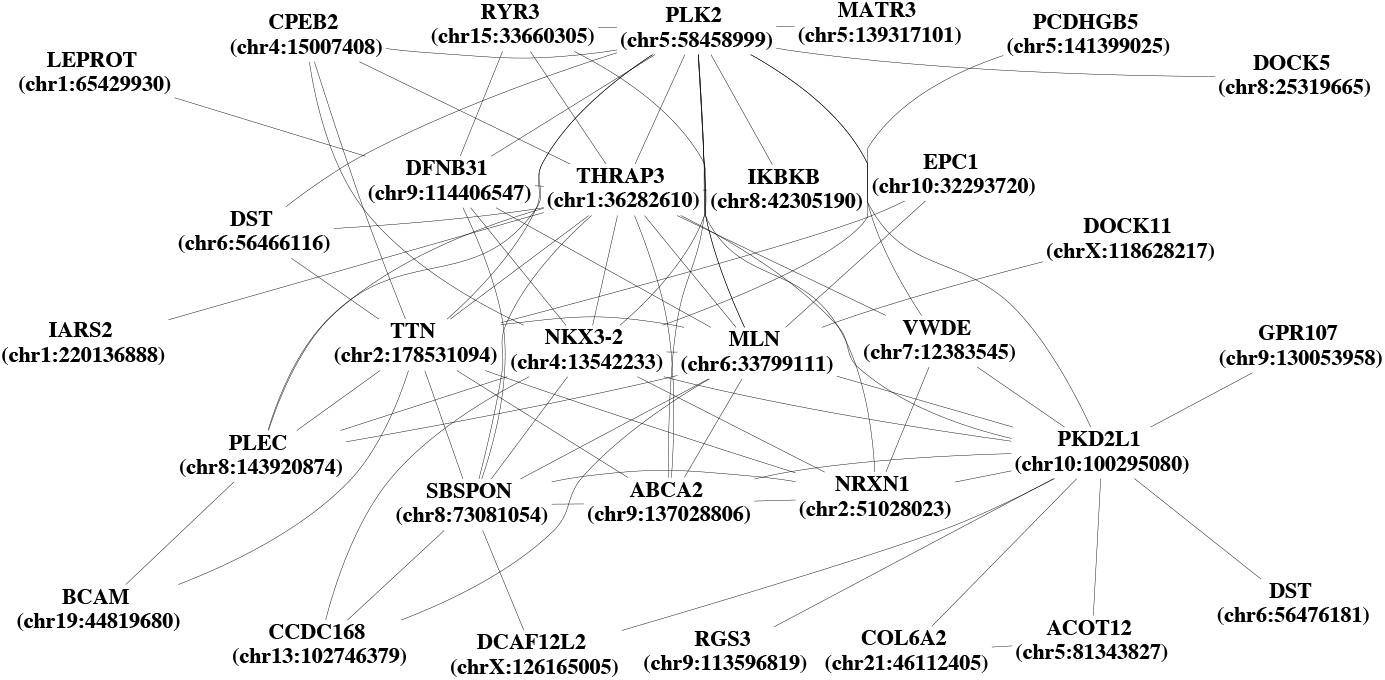
Inferred mutation co-occurence network across Patient 6from Model hGP.

